# Efficient Exact Inference for Dynamical Systems with Noisy Measurements using Sequential Approximate Bayesian Computation

**DOI:** 10.1101/2020.01.30.927004

**Authors:** Yannik Schälte, Jan Hasenauer

## Abstract

**Motivation:** Approximate Bayesian Computation (ABC) is an increasingly popular method for likelihood-free parameter inference in systems biology and other fields of research, since it allows analysing complex stochastic models. However, the introduced approximation error is often not clear. It has been shown that ABC actually gives exact inference under the implicit assumption of a measurement noise model. Noise being common in biological systems, it is intriguing to exploit this insight. But this is difficult in practice, since ABC is in general highly computationally demanding. Thus, the question we want to answer here is how to efficiently account for measurement noise in ABC.

**Results:** We illustrate exemplarily how ABC yields erroneous parameter estimates when neglecting measurement noise. Then, we discuss practical ways of correctly including the measurement noise in the analysis. We present an efficient adaptive sequential importance sampling based algorithm applicable to various model types and noise models. We test and compare it on several models, including ordinary and stochastic differential equations, Markov jump processes, and stochastically interacting agents, and noise models including normal, Laplace, and Poisson noise. We conclude that the proposed algorithm could improve the accuracy of parameter estimates for a broad spectrum of applications.

**Availability:** The developed algorithms are made publicly available as part of the open-source python toolbox pyABC (https://github.com/icb-dcm/pyabc).

**Supplementary information:** Supplementary information is available at *bioRxiv* online. Supplementary code and data are available online at http://doi.org/10.5281/zenodo.3631120.

## 1 Introduction

Mathematical models have become an essential tool in many research areas to describe and analyse dynamical systems, allowing to unravel and understand underlying mechanisms. In order to make quantitative predictions and test hypotheses, unknown parameters need to be estimated and parameter and prediction uncertainties need to be assessed.

This is frequently done in a Bayesian framework, where prior information and beliefs about model parameters are updated by the likelihood of observing data under a given model parameterization, yielding by Bayes’ Theorem the posterior distribution of the parameters given the data. Many established parameter estimation methods, including optimization (Banga, 2008), profile calculation (Raue *et al*., 2009) and standard Monte Carlo sampling (Hines, 2015), require access to at least the non-normalized posterior. However, as models get more complex and stochastic, the likelihood function can become analytically or numerically intractable (Jagiella *et al*., 2017). Examples of such models include Markov processes, stochastic differential equations, and stochastically interacting agents. In systems biology, such models are used to realistically describe e.g. gene expression, signal transduction, and multi-cellular systems (e.g. Lenive *et al*. (2016); Picchini (2014); Imle *et al*. (2019)).

Likelihood-free inference methods have therefore recently gained interest, including among others Approximate Bayesian Computation (ABC) (Beaumont *et al*., 2002; Sisson *et al*., 2018b), indirect inference (Gourieroux *et al*., 1993; Drovandi, 2018), synthetic likelihoods (Wood, 2010; Price *et al*., 2018), and particle Markov Chain Monte Carlo (Andrieu *et al*., 2010). In particular, ABC has become increasingly popular in various research areas due to its simplicity, scalability, and its broad applicability. In a nutshell, in ABC the evaluation of the likelihood function is circumvented by simulating data for given parameters, and then accepting the parameters if the simulated and observed data are sufficiently similar. ABC is frequently combined with a sequential Monte Carlo scheme (ABC-SMC) (Del Moral *et al*., 2006; Sisson *et al*., 2007), which allows for an iterative reduction of the acceptance threshold, improves acceptance rates by sequential importance sampling, and can exploit parallel infrastructure well.

Observed data are generally corrupted by noise, resulting from unavoidable inaccuracies in the measurement process. In likelihood-based inference, it has been widely adopted to include noise models in the likelihood function (Raue *et al*., 2013). Contrarily, in likelihood-free methods, particularly ABC, it is easy to disregard any noise due to the unnecessity of even formulating a likelihood and the various inherent approximation levels, so that error sources can be difficult to pinpoint from the result. In the past, it has repeatedly not been included in ABC analyses (Toni *et al*., 2009; Lenive *et al*., 2016; Jagiella *et al*., 2017; Imle *et al*., 2019; Eriksson et al.,2019). Asymptotic unbiasedness of ABC is however granted only if the data-generation process is perfectly reproduced. Omitting the measurement noise can lead to substantially wrong parameter estimates, regarding in particular uncertainty (Frazier *et al*., 2020).

The problem is illustrated in Figure 1 on an ordinary differential equation model of a conversion reaction, with one unknown parameter *θ*_1_. Synthetic data *D* were generated by adding normal noise to the model simulation *y*(*θ*_true_) (Figure 1A). Three different ABC-SMC analyses were performed: Using the noise-free model *y* together with an *ℓ*_1_ (I) or *ℓ*_2_ (II) distance, and, to account for noise, randomizing the model output by a corresponding normal random variable (III). Usually, in ABC we would hope to decrease the acceptance threshold *ε* asymptotically to 0. For (I) and (II), this was however not possible (Figure 1B). The thresholds converged to some positive values, which in addition differ for the *ℓ*_1_ and *ℓ*_2_ distance. This mirrors in the inferred posterior distributions (Figure 1C), where (I) and (II) converge to point estimates, which can be linked to maximum likelihood estimates under the assumption of normal (I) or Laplace (II) noise. This is clearly not the result one would hope for in a Bayesian analysis. In comparison, (III) yields a good approximation of the true posterior. In the Supplementary Information, Section 1.5 we discuss the problem of model error in ABC from a theoretical perspective, also explaining what happens in the above analyses. Further, in the Supplementary Information, Section 6 we illustrate on some more examples, including stochastic models, how ignoring measurement noise can lead to wrong parameter estimates. In practice, errors in the ABC parameter estimates resulting from model error can be hard to detect, so it is important to correctly account for measurement noise.

**Figure 1:**
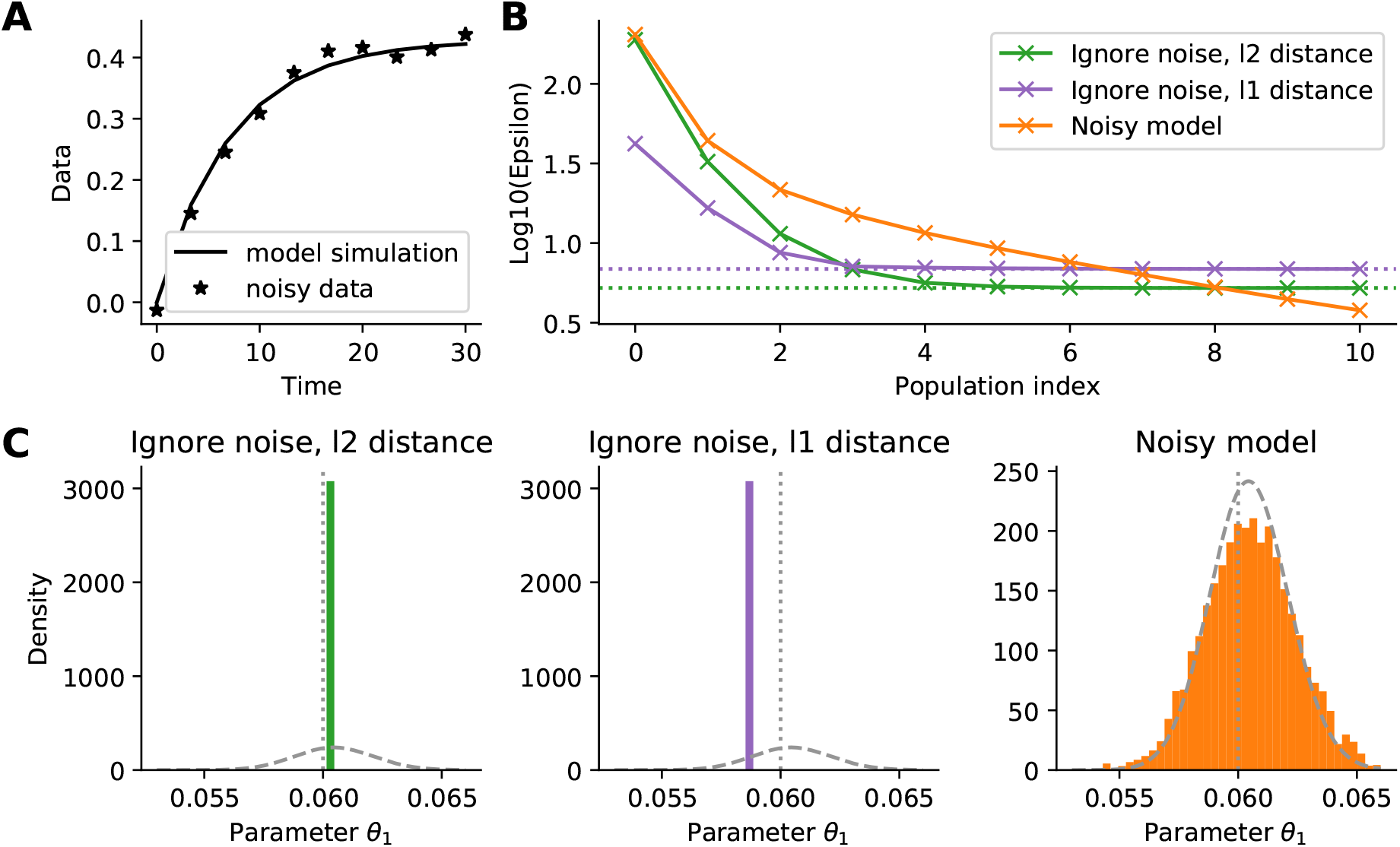
Illustrative conversion reaction ODE example. (A) The employed data. (B) Acceptance thresholds over sequential ABC-SMC iterations for 3 estimation methods: Using a non-noisy model and an *l*_2_ or *l*_1_ distance in the ABC acceptance step, and using a noisy model (and an *l*_2_ distance). The minimum obtainable values for the respective distances are indicated by dashed lines. (C) Histograms of the corresponding ABC posterior approximations after the last iterations. The true posteriors are indicated by dashed, the true parameter values by dotted lines.

In this manuscript, we discuss ways of correctly addressing measurement noise: Either the model output can be randomized, or the ABC acceptance step can be modified in accordance with the noise model. The latter method builds on the insight by Wilkinson (2013) that ABC can be considered as giving exact inference from the original model with an additional error term induced by the acceptance step. Introduced by Wilkinson (2013) for rejection and Markov Chain Monte Carlo (ABC-MCMC) samplers and used by van der Vaart *et al*. (2018) in the case of replicate measurements, the approach was extended by Daly *et al*. (2017) to ABC-SMC samplers,presenting two algorithms confined to the situation of additive independent normal noise, and relying on certain tuning parameters. Here, we extend the existing ideas by presenting an ABC-SMC based algorithm applicable to various model types and noise models. Further, we develop robust approaches tackling several aspects of such an algorithm, like initialization and step size selection, and include ideas from rejection control importance sampling (Liu *et al*., 1998; Sisson and Fan, 2018), which make our algorithm robust and self-tuned and thus widely applicable. We test and compare it on various models including ordinary and stochastic differential equations, discrete Markov jump processes, and agent-based models, and noise models including normal, Laplace, and Poisson noise.

## 2 Methods

### 2.1 Basics of ABC-SMC

The goal of Bayesian inference is to infer a posterior distribution *π*(*θ*|*D*) ∝ *p*(*D*|*θ*)*π*(*θ*) over parameters 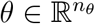 given observed data 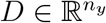, where *π*(*θ*) denotes the prior density on the parameters encoding information and beliefs before observing the data, whereas the likelihood *p*(*y*|*θ*) is the probability density of data *y* given model parameters *θ*. ABC deals with the situation that we have a generative model from which we can simulate data *y* ~ *p*(*y*|*θ*), but evaluating the likelihood is infeasible. Then, classical ABC comprises the following 3 steps:

1. sample parameters *θ* ~ *π*(*θ*),
2. simulate data *y* ~ *p*(*y*|*θ*),
3. accept *θ* if *d*(*y*, *D*) ≤ *ε*,

for some distance function 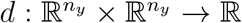 and acceptance threshold *ε*. This is repeated until sufficiently many, say *N*, parameters have been accepted. Accepted particles constitute a sample from the approximate posterior distribution

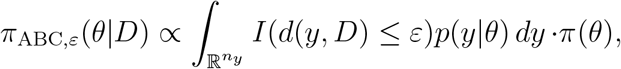

where *I* denotes the indicator function, also referred to as *uniform kernel* in this context.

Due to the curse of dimensionality, ABC usually works not directly on the data, but employs low dimensional summary statistics 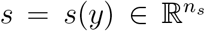 (Fearnhead and Prangle, 2012). Here, we abstract from this to simplify the notation. The summary statistics may be assumed to be already incorporated in *y*.

It can be shown that for *ε* → 0, *π*_ABC,*ε*_(*θ*|*D*) → *π*(*θ*|*D*) under some mild assumptions (e.g. Prangle *et al*. (2017), and Supplementary Information, Section 1.4), however there is a trade-off between decreasing the approximation error induced by *ε* and maintaining high acceptance rates. To tackle both problems, ABC is frequently combined with a Sequential Monte Carlo (SMC) scheme. Here, we present a scheme based on Toni *et al*. (2009); Beaumont *et al*. (2009). In ABC-SMC, a population of parameters 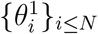 is initially sampled from the prior and propagated through a sequence of intermediate distributions *π*(*θ*|*d*(*y*\*, *D*) ≤ *ε_t_*), *t* = 1,…,*n_t_*, using importance sampling, with the proposal distribution *g_t_*(*θ*) based on the previous iteration’s accepted particles. The tolerances *ε*_1_ > … > *εn_t_* ≥ 0 are chosen to yield a gradually better approximation of the posterior distribution, while maintaining high acceptance rates. The steps are summarized in Algorithm 1.

#### Algorithm 1 ABC-SMC algorithm

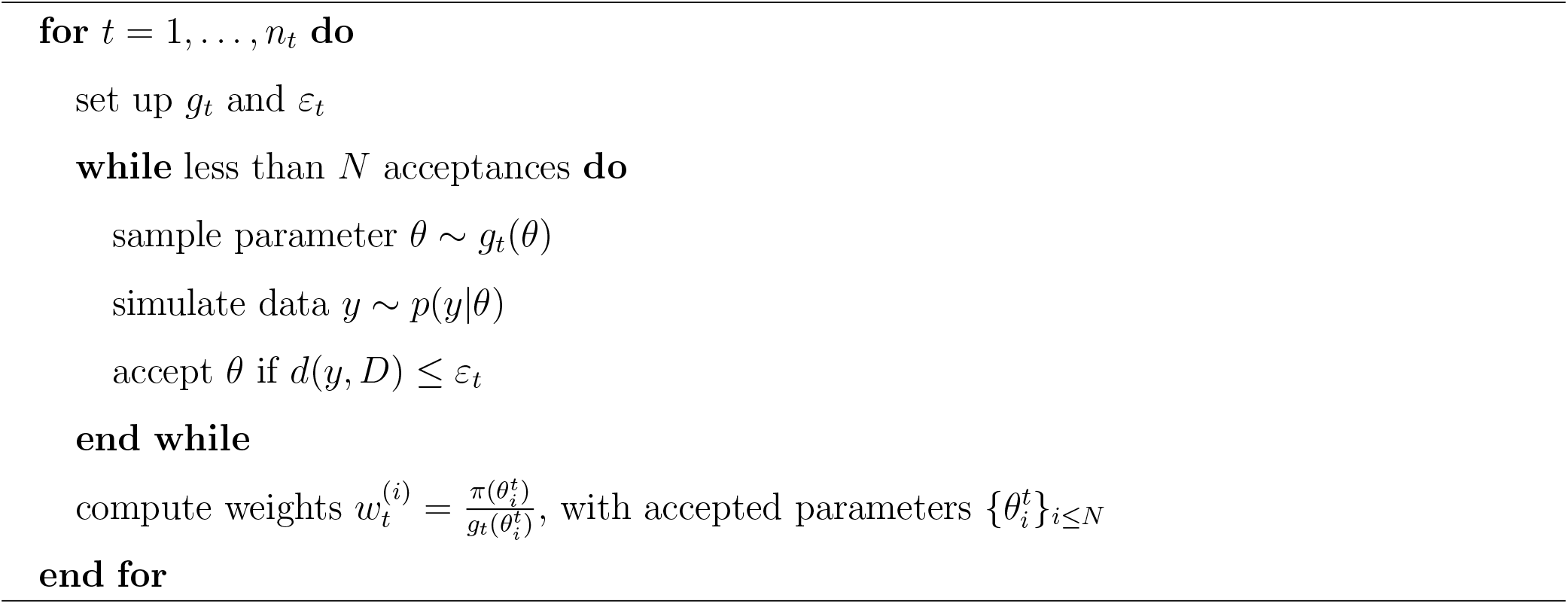

Here, the proposal distribution in iteration *t* is

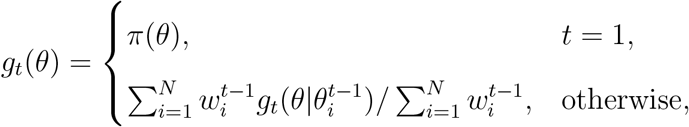

with 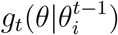 local perturbation kernels, 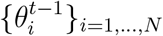 the accepted parameters in the previous iteration, and 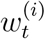 importance weights (Klinger and Hasenauer, 2017). The output of ABC-SMC is a population of weighted parameters 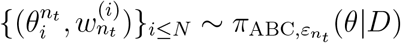.

### 2.2 The problem of measurement noise in ABC

In ABC, the model *p*(*y*|*θ*) does often not account for measurement noise. This effectively assumes perfect measurements, which is in practice hardly the case. In the following, we assume that the data *D* are noisy and thus a realization of a distribution *q*(*ȳ*|*θ*) which includes the noise, but that the model *p*(*y*|*θ*) does not do so. Further, we can write (*ȳ*, *y*) ~ *π*(*ȳ*|*y*, *θ*)*p*(*y*|*θ*), so that

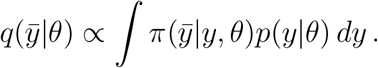

The interpretation of *π*(*ȳ*|*y*, *θ*) is that of a parameterized noise model of observing data *ȳ* under noise-free model output *y* and parameters *θ*. Thus, the noise model relates observables that assume perfect measurements to practically obtained noisy data. We call

- *π*(*ȳ*|*y*, θ) the *noise model*,
- *p*(*y*|*θ*) the *model likelihood*, and
- *q*(*ȳ*|*θ*) the *full likelihood*.

The noise model *π*(*ȳ*|*y*, *θ*) is usually simple, e.g. a normal or a Laplace distribution. Hence, we consider the case that the noise model can be evaluated, while we are only able to sample from, but not evaluate, the model likelihood.

The use of the likelihood *p* as above would imply that inference for the wrong model is performed. The goal is now to infer the corrected posterior *π*(*θ*|*D*) ∝ *q*(*D*|*θ*)*π*(*θ*).

### 2.3 Approaches to account for noise

How can we tackle in ABC the discrepancy between model output *y* and noisy data *D*? Principally, three different approaches have been in use, which are visualized in Figure 2.

**Figure 2:**
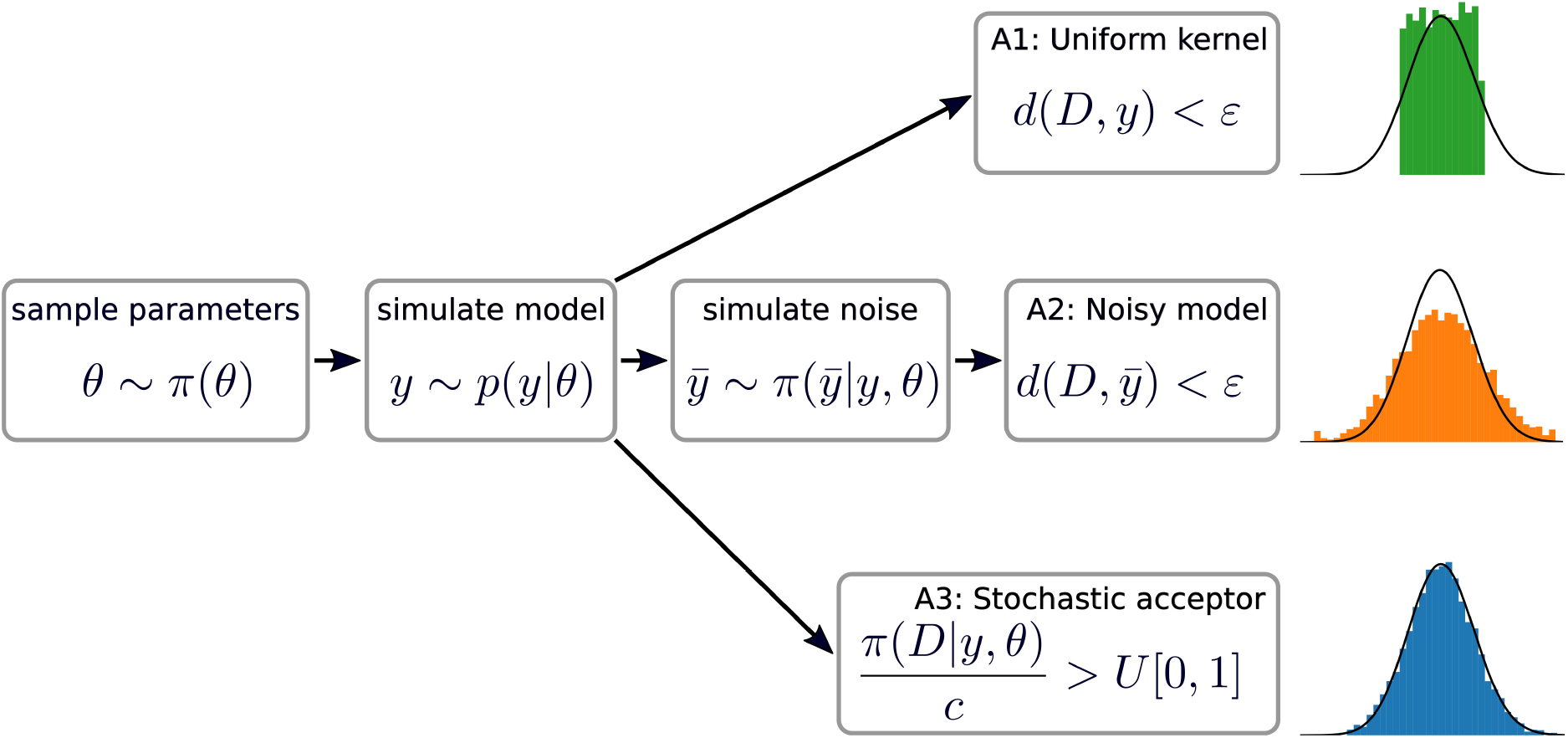
The different conceivable ways of accounting for noise. (A1): Using a uniform kernel with *ε* ≫ 0. The posterior is visibly off. (A2): Adding random measurement noise to the model output. The posterior still has a slightly higher variance. (A3): Modifying the acceptance kernel. The posterior matches the true distribution accurately.

#### Using an appropriate uniform kernel

The first approach (A1) is to employ a uniform kernel, with the distance metric *d* and the acceptance threshold *ε* chosen such that the resulting acceptance kernel is similar to the underlying noise distribution (e.g. in Toni *et al*. (2009); Daly *et al*. (2017)). An advantage of this approach is its computational efficiency, since acceptance is deterministic (Sisson *et al*., 2018a). Further, it is easy to apply in practice since it only uses standard ABC methods available in most software tools. However, a major concern is that this approach effectively assumes a uniform noise distribution (see Theorem 1, Section 2.4), which in practice hardly applies. In addition, the exact choice of *ε* is ambiguous. E.g., one can fix the kernel variance, or set it to the expected value of the distance function at the true noise-free model value. Doing so requires knowledge of underlying noise parameters, or even of *y*. This information is in practice not available. Here, we only mention this approach for completeness, and focus on (asymptotically) exact methods.

#### Randomizing the model output

The second approach (A2) is to modify the forward model simulation to account for measurement noise, i.e. to randomize the model output *y* → *ȳ* ~ *π*(*ȳ*|*y*, *θ*) (e.g. in Toni and Stumpf (2010)), thus replacing *p* by *q*. An advantage of this method is that it is again easy to apply, requiring only a basic ABC implementation. Further, if the noise model depends on unknown parameters, these can in theory be included in the overall parameter vector *θ* and estimated along the way. Also, this approach is in particular applicable to “black box” models where the noise cannot be separated. A major concern with this method is however that the randomness in the simulation of noise can lead to low acceptance rates. Further, the comparison of simulated and observed data still requires a standard acceptance kernel with a non-trivial threshold. A2 is asymptotically exact as *ε* → 0, however in practice a small approximation error will remain, which can be hard to quantify.

#### Modifying the acceptance step

The third approach (A3) is to keep the non-noisy model *p*, but to modify the acceptance criterion: Based on the insight by Wilkinson (2013) that “ABC gives exact inference, but for the wrong model”, we modify the acceptance step from Section 2.1 to 3. accept *θ* with probability 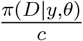, with a normalization constant *c* ≥ max_*y*,*θ*_ *π*(*D*|*y*, *θ*). This step can be implemented by sampling *u* ~ *U*[0, 1] and accepting if 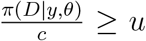. Theorem 1 (Section 2.4) tells us that this indeed gives exact inference from the true posterior. It is thus the non-degenerate noise model that allows us to perform likelihood-free inference in an exact manner (i.e., up to Monte Carlo errors), still without evaluating the full likelihood. Only for deterministic models do we have the full likelihood, such that this approach is equivalent to likelihood-based sampling techniques. In Wilkinson (2013), this idea has been integrated in an ABC-rejection and an ABC-MCMC algorithm. In Daly *et al*. (2017), two sequential implementations for Gaussian noise are introduced. Building on both works, we present in the following a self-tuned sequential algorithm applicable to a broad class of noise models.

### 2.4 ABC gives exact inference under the assumption of model error

We use the following:

#### Theorem 1

(Exact noisy ABC). *Consider a prior density π*(*θ*), *a model likelihood p*(*y*|*θ*), *a noise model π*(*ȳ*|*y*, *θ*), *and assume D* ~ *q*(*ȳ*|*θ*) ∝ ∫*π*(*ȳ*|*y*, *θ*)*p*(*y*|*θ*) *dy*. *Then ABC with acceptance probability π*(*D*|*y*, *θ*)/*c with c* ≥ sup_*y*,*θ*_ *π*(*D*|*y*, *θ*) *targets the correct posterior distribution π*(*θ*|*D*) ∝ *q*(*D*|*θ*)*π*(*θ*).

*Proof*. See the Supplementary Information, Section 2.

This extends the work of Wilkinson (2013) to generic noise distributions and allows the noise model to be parameter-dependent. The latter facilitates, e.g., the estimation of unknown scale terms such as the standard deviation for normally distributed measurement noise. This result can also be interpreted in the context of general ABC acceptance kernels, see the Supplementary Information, Section 1.3 and Remark 4.

### 2.5 Towards an efficient exact sequential ABC sampler

To increase efficiency, we want to integrate the exact sampler A3 with an SMC scheme. To do so, we need to replace the gradual decrease of the acceptance threshold *ε*. The basic idea we employ is motivated by parallel tempering in MCMC (Earl and Deem, 2005), namely to temper the acceptance kernel to mediate from prior to posterior. That is, we introduce temperatures *T*_1_ > … > *T_n_t__* = 1, and in iteration *t* modify the acceptance step to 3. accept *θ* with probability 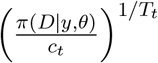.

This way, we sample from the distribution

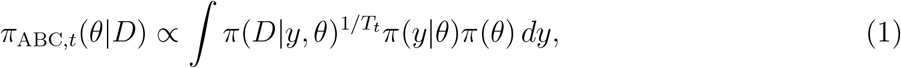

using importance sampling, where *T_n_t__* = 1 yields a sample from the correct posterior. Note that tempering is applicable to any noise model and will yield higher acceptance rates. An exception is the uniform distribution, for which the acceptance rate will remain unchanged, but this noise model can be dealt with by standard ABC already. In the following, we propose approaches to select the normalization constant *c*, the temperature schedule, and the initial temperature.

### 2.6 Selection of the normalization constant

A problem persistent in the approaches by Wilkinson (2013) and Daly *et al*. (2017) was the choice of normalization constant *c*. A trivial choice for this is the highest mode of the noise distribution, which is for common noise models assumed at *y* = *D*. Yet, in practice it is often unlikely or impossible for the model to exactly replicate the measured data, yielding unnecessarily small acceptance rates. Thus, of interest is the point 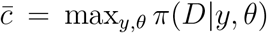 such that *y* is realizable under the model *p*(*y*|*θ*). For deterministic models, this point is the maximum likelihood value and can be computed by optimization. For stochastic models, it is in general unknown. Daly *et al*.(2017) argue that too small a *c* leads to a decapitation and uniformization of the noise distribution around the maximum likelihood value. Due to the inability to find good values for *c*, there the here employed ABC-SMC sampler based on Toni *et al*. (2009) was disregarded in favor of a sampler based on Del Moral *et al*. (2012), although the former had shown superior accuracy. We can however solve both the problems of too low acceptance rates and of the decapitation of the noise model by correcting for that error as follows, based on ideas from rejection control importance sampling (RCIS, Sisson and Fan (2018)):

#### Theorem 2

(Importance weighted acceptance). *Let c_t_* > 0 *arbitrary. If we change the acceptance step to* 3. accept with probability min 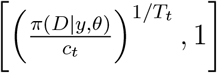 *and modify the importance weights to be*

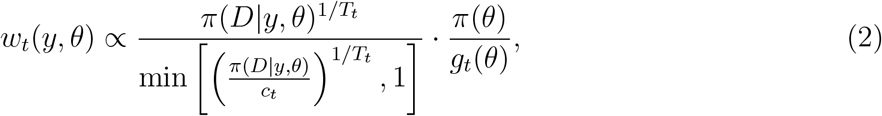

*the weighted samples* (*θ, w*(*θ*)) *target distribution* (1).

*Proof*. See the Supplementary Information, Section 3.3.

This means that we can, for arbitrary *c*, correct for accepting from the decapitated noise distribution by modifying the acceptance weights. Of course, a smaller *c* leads to higher acceptance rates, but also to an increase in the Monte Carlo error by weight degeneration. Therefore, it must be chosen carefully. In a sequential approach, it is straightforward to iteratively update *c* by taking into account previously observed values. In this study, we by default set it to the maximum of the values found in previous iterations. When acceptance rates turn out too low, we set it to *β*^−*T*^*C*, where *c* is the maximum found value, and *β* ≥ 1 increases acceptance rates roughly by that factor in the next iteration. Other schemes, e.g. based on quantiles, are possible. Before iteration 1, we draw a calibration sample from the prior.

### 2.7 Selection of the temperatures

A proper temperature scheme has to balance information gain and acceptance rate. In general, the overall required number of simulations depends both on the number of intermediate populations, and the difficulty of jumping between subsequent distributions. In the following, we propose two schemes based on different criteria.

#### Acceptance rate scheme

The idea of this scheme is to match a specified target acceptance rate, i.e. to choose *T* = *T_t_* such that the expected acceptance rate

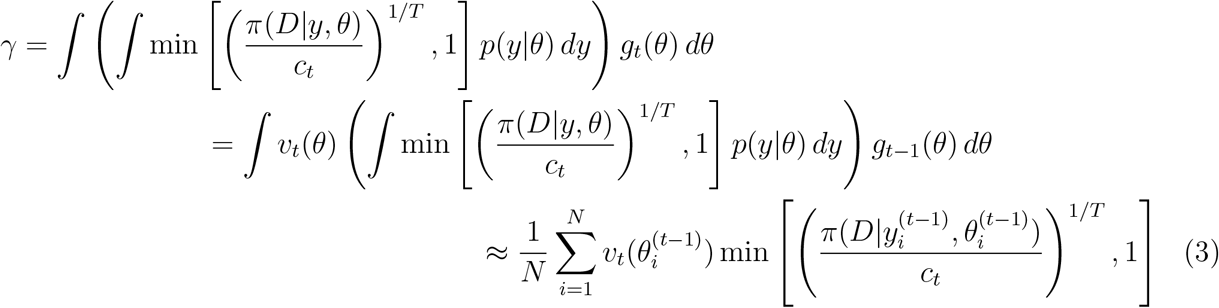

matches a specified target rate. Here, in the second line we employ importance sampling from the previous proposal distribution *g*_*t*−1_ with corresponding Radon-Nikodym derivatives *v_t_*(*θ*) = *g_t_*(*θ*)/*g*_*t*−1_(*θ*). This is because in the third line, we approximate this integral via a Monte Carlo sample, using all parameters sampled in the previous iteration. This must include rejected particles to avoid a bias. The inner integral is approximated by the corresponding single simulation.

Matching *γ* ≈ *γ*_target_ is a one-dimensional bounded optimization problem which can be efficiently solved. Compared to the overall run time of ABC analyses, we found the computation time to be negligible. While this scheme provides only a rough estimate of the expected acceptance rate, it proved sufficient for our purpose.

Assuming convergence *c_t_* → *c*_∞_ and *g_t_* → *g*_∞_ similar in shape to the true posterior distribution, it is to be expected that a such proposed *T* converges to a value *T*_∞_ > 1 in general. Therefore, the acceptance rate scheme is rather intended as a scheme for the first few iterations and needs to be backed up with an additional scheme, e.g. the one following, that ensures *T* ↘ 1.

#### Exponential decay scheme

In standard likelihood-based parallel tempering MCMC, empirically a geometric progression, i.e. a scheme with fixed temperature ratios, has shown to yield roughly equal probabilities for swaps between adjacent temperatures (Sugita *et al*., 2000). This finding can be theoretically justified (Predescu *et al*., 2004). Since a similar approach was recently successfully applied in an ABC-SMC setting by Daly *et al*. (2017), we used a geometric progression here as well. We specified a fixed ratio *α* ∈(0,1) such that *T*_*t*+1_ = *αT_t_*.

The effective next temperature was then set to the minimum of the temperatures proposed by the acceptance rate scheme and the exponential decay scheme.

#### Find a good initial temperature

Unanswered has remained the question of how to choose the initial temperature. It should be low enough to avoid simply sampling from the prior without information gain, but have reasonable acceptance rates. Thus it is a crucial tuning parameter. Here, we propose a widely applicable self-tuned mechanism based on the above acceptance rate scheme: The acceptance rate scheme can be applied to find *T*_1_ if we set *g*_1_(*θ*) = *g*_0_(*θ*) = *π*(*θ*) and generate a calibration sample 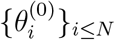 from the prior, the same we use to find the initial normalization *c*.

For brevity, we denote by ASSA in the following the here proposed exact ABC-SMC sampler with an adaptive sequential stochastic acceptor, i.e. with *c* set to the previously observed highest value with weight correction, and the two temperature selection schemes, with target acceptance rate *γ*_target_ = 0.3, and *α* = 0.5 in the exponential decay scheme.

### 2.8 Implementation

We implemented all the algorithms in the open-source python toolbox pyABC (https://github.com/icb-dcm/pyabc, Klinger *et al*. (2018)), which offers a state-of-the-art implementation of ABC-SMC. We put emphasis on an easy-to-use modular implementation, such that it is straightforward to customize the analysis pipeline. To ensure numerical stability, critical operations were performed in log-space. For further details see the Supplementary Information, Section 5. Jupyter notebooks illustrating how to use the algorithms have been included in the pyABC online documentation. The complete data and code are available on zenodo (http://doi.org/10.5281/zenodo.3631120).

## 3 Results

To study the properties of ASSA, and compare it to alternative approaches, we consider six models.

### 3.1 Model description

The models cover various modeling formalisms, including ordinary differential equations (ODE), stochastic differential equations (SDE), Markov jump processes (MJP), and agent-based models (ABM), as well as various noise models, including normal, Laplace, and Poisson noise.

Models M1 and M2 are ODE models of a conversion reaction 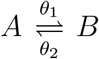, a typical building block in many biological systems. We estimated the reaction rate coefficients *θ*_1_, *θ*_2_, assuming only species *A* to be measured. In model M1, we employed independent additive normal noise, a common assumption in systems biology. In model M2, we instead employed Laplace noise, which is frequently used when the data are prone to outliers (Maier *et al*., 2017). Further, M1 was varied in multiple regards to investigate various features of the algorithm.

Model M3 is an SDE model of intrinsic ion channel noise in Hodgkin-Huxley neurons based on Goldwyn *et al*. (2011). We estimated the parameters dc describing the input current, and the square root of the membrane area membrane_dim. We assumed measurements to be available for the fraction of open potassium channels *K*, and employed an additive normal noise model to describe inaccuracies in the measurement process.

Model M4 describes the process of mRNA synthesis and decay, 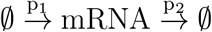, we estimated the transcription rate constant *p*_1_ and the decay rate constant *p*_2_. To capture the intrinsic stochasticity of this process at low copy numbers, we sampled from the chemical master equation using the Gillespie algorithm (Gillespie, 1977). We assumed the counts of mRNA molecules from a single cell to be available at discrete time points. Data of this kind can be obtained e.g. by fluorescence microscopy. We employed a Poisson noise model, which is frequently used for regression of count data (Coxe *et al*., 2009).

Model M5 is an ODE model by Boehm *et al*. (2014) describing the homo- and heterodimerization of the transcription factors STAT5A and STAT5B, for which three types of data with 16 measurements each are available. In the original publication, additive normal measurement noise was assumed, and optimal parameter point estimates obtained using optimization. We estimated 11 logarithmically scaled parameters of this model, including three standard deviations of the normal noise model, one for each data type.

Model M6 is a multi-scale ABM model of spheroid tumor growth on a two-dimensional plane, as described in Jagiella *et al*. (2017). Single cells were modeled as stochastically interacting agents, coupled to the dynamics of extracellular substances modeled via partial differential equations. For the three data types generated by this model, we assumed normal noise models of differing variance. The model has seven unknown parameters. The simulation of this ABM model is computationally relatively demanding, and a single forward simulation on the employed hardware took already about 20 seconds. As often on the order of 1e5 to 1e8 forward simulations are required for inference, the overall computation time was on the order of thousands of core hours.

For models M1-4 and M6, we created artificial data by simulating the model and then “noisifying” the simulations by sampling from the respective noise distribution. Model M5 is based on real data without known ground truth but reported literature values. For all models, we used uniform parameter priors over suitable ranges. A summary of the model properties is provided in Table 1, and further details, including a visualization of the employed data, can be found in the Supplementary Information, Section 7.

**Table 1:**
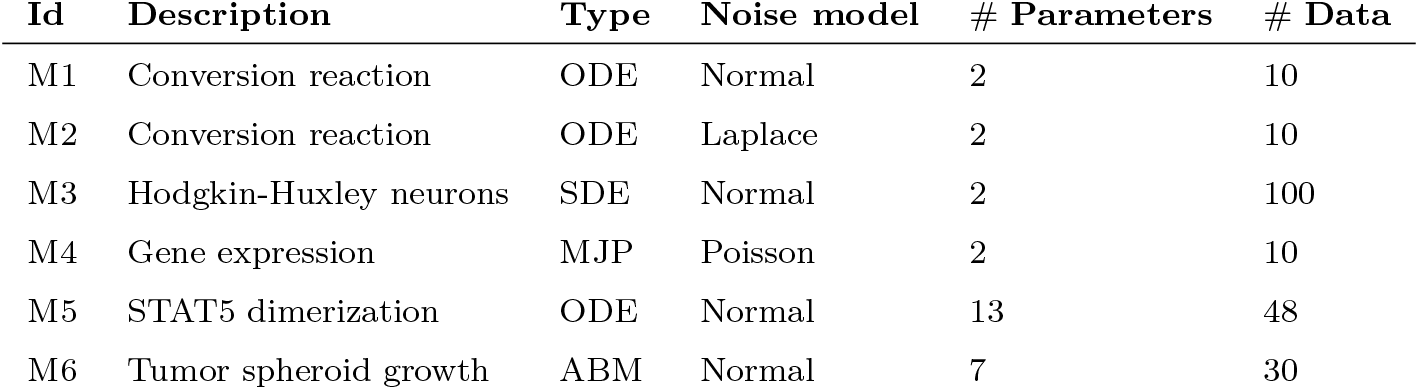
Main properties of the considered models.

| **Id** | **Description** | **Type** | **Noise model** | **# Parameters** | **# Data** |
| --- | --- | --- | --- | --- | --- |
| M1 | Conversion reaction | ODE | Normal | 2 | 10 |
| M2 | Conversion reaction | ODE | Laplace | 2 | 10 |
| M3 | Hodgkin-Huxley neurons | SDE | Normal | 2 | 100 |
| M4 | Gene expression | MJP | Poisson | 2 | 10 |
| M5 | STAT5 dimerization | ODE | Normal | 13 | 48 |
| M6 | Tumor spheroid growth | ABM | Normal | 7 | 30 |

For parameter estimation, we used pyABC with a multivariate normal kernel with adaptive covariance matrix as proposal distribution between populations, and a median strategy to update the *ε* threshold under a uniform acceptance kernel (Klinger and Hasenauer, 2017). If not mentioned otherwise, we used a population size of *N* = 1000.

### 3.2 Reweighting reliably corrects for bias

As the RCIS reweighting derived in Section 2.6 should allow for exact inference independent of the normalization constant *c*, we compared the resulting sample distribution with the ground truth. Therefore, we performed sampling for different values of *c*, for model M1 confined to one estimated parameter (Figure 3A). The theoretical maximum value of the likelihood of the data *D* under the model was determined by multi-start local optimization, yielding 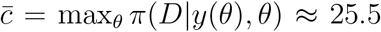. Here, we used *c* ∈ {20, 23, 25}. As already pointed out in Daly *et al*. (2017), this leads to a decapitation of the posterior distribution, i.e. it flattens out at values of high probability. The difference to the true posterior became smaller the larger *c*. However, if we corrected for the bias introduced by the too low normalization according to (2), we obtained, even for the lowest *c* = 20, a posterior that well matched the true distribution.

**Figure 3:**
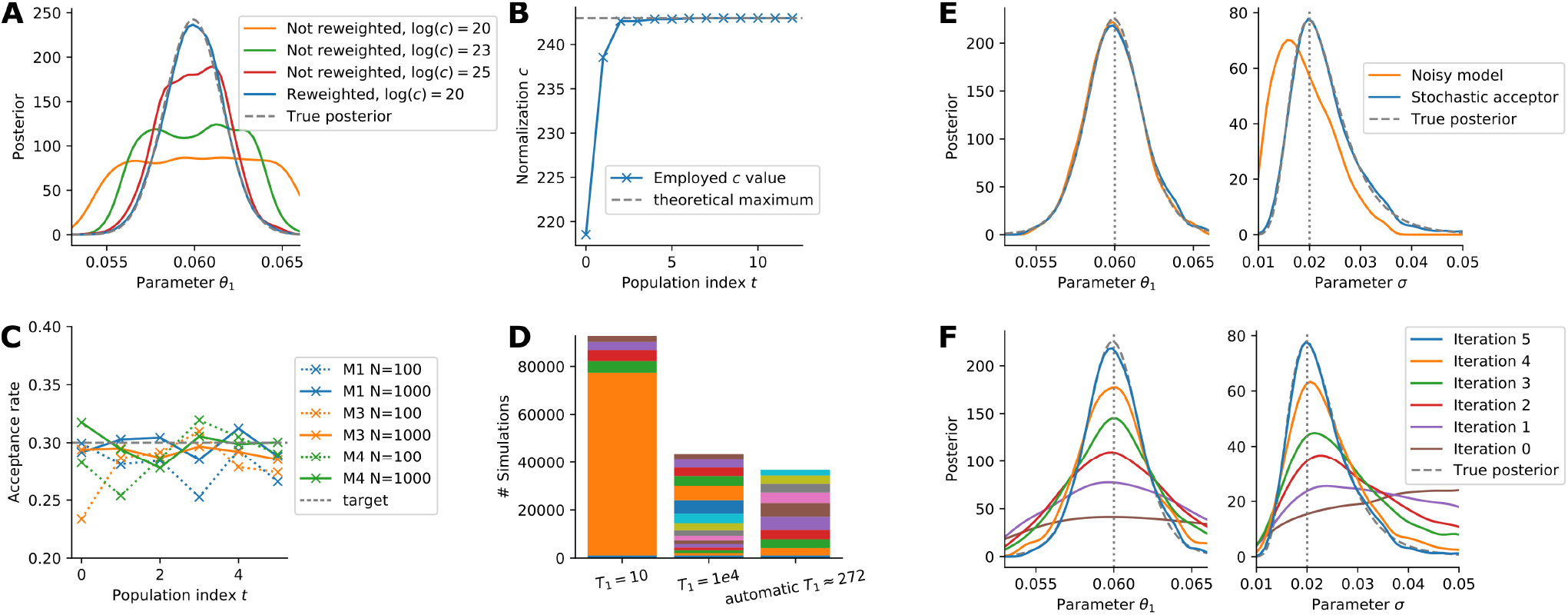
Evaluation of the properties of the components of the proposed sampling scheme. (A) Kernel density estimates (KDE) of four posterior estimates generated for model M1 with one unknown parameter and 10 data points, with different normalization constants *c*, only once correcting for the acceptance bias by reweighting. (B) Normalization constant *c* over the iterations for inference for model M1 with 100 data points. (C) Acceptance rate in sampling runs for models M1, M3 and M4 for two population sizes *N*. Only the acceptance rate criterion was used for temperature scheduling. (D) Total number of simulations in runs for model M1 for different initial temperatures *T*_1_. The temperature updates were performed using only the exponential decay scheme. The colors indicate individual iterations, starting at the bottom with the simulations spent in the calibration iteration, and then from *t* = 1 upwards. (E,F) KDE of simulations obtained for model M1 with one unknown dynamic parameter, and also estimating the normal noise variance. (E) Comparison of a run using the stochastic acceptor, and a run using a noisy model output. (F) Posterior estimates over the sequential iterations using the stochastic acceptor.

Employing the proposed strategy to automatically update *c* after each iteration to the highest value so far, we observed for all test models that the value of *c* converged over time, larger jumps taking place only in the first few iterations. E.g. for the 2-parametric model M1 with 100 data points (Figure 3B), *c* converged to the theoretical minimum upper bound 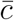.

### 3.3 Acceptance rate prediction works reliably

Next, we analysed the performance of the acceptance rate prediction introduced in Section 2.7 as a means to choose temperatures. We ran six iterations for each of the models M1-3 and found that the deviations to the target value of *γ*_target_ = 0.3 are acceptable (Figure 3C). The fit was already good at *t* = 1, thus allowing to find appropriate initial temperatures. As expected, the fluctuations decreased for a higher population size. This shows that approximation (3) is sufficient for the temperature adaptation.

While the acceptance rate criterion provided a means to adapt the temperature in early iterations, the proposed temperatures *T* did in most cases not converge to *T* = 1, at which we have exact inference. This was to be expected, as explained in Section 2.7, and necessitates the presence of a secondary scheme ensuring *T* ↘ 1, for example the exponential decay scheme. We observed that the acceptance rate criterion reliably proposed good initial temperatures and allowed for major temperature jumps in the first iterations (accelerating convergence), while in later iterations the exponential decay scheme took over (e.g. Supplementary Information, Figure S10).

To illustrate the importance of a proper selection of the initial temperature, we fixed it for model M1 to different values (Figure 3D), employing only the exponential decay scheme with a fixed number of iterations. Too small values of the initial temperature led to many simulations being necessary in the initial iteration, sometimes even more than for the entire analysis using the self-tuned initial temperature. Too high values of the initial temperature yielded little information gain and resulted in a waste of computation time in the first iterations.

### 3.4 Approach allows to estimate noise parameters

In Theorem 1, we allowed the noise likelihood *π*(*ȳ*|*y*, *θ*) to be parameter-dependent for the stochastic acceptor. To test the validity of this, we employed model M1 and estimated the standard deviation *σ* of the normal noise model along with the rate constant *θ*_1_ (Figure 3E). Indeed, the posterior distributions of both parameters obtained using the proposed algorithm match the ground truth. In theory, also employing approach A2, i.e. a noisy model, should approximately allow to estimate noise parameters. However, even after 6*e*6 simulations, compared to 1*e*5 for the stochastic acceptor, the quality of the estimated posterior distribution for the noise parameter was considerably worse. Meanwhile, parameter *θ*_1_ was well estimated.

The sequential improvement of the posterior distribution over the ABC-SMC iterations for the same model (Figure 3F) indicates that the temperature update scheme suggests steps with an appropriate information gain.

### 3.5 Applicable to various model types and noise models

To evaluate how our proposed ASSA sampler performs on different types of dynamical models *p*(*y*|*θ*) and noise models *π*(*ȳ*|*y*, *θ*), we studied models M1-4. Further, we compared ASSA to alternative approaches: Firstly, we employed it with only one iteration, *n_t_* = 1, giving exact ABC-rejection. Secondly, we set the normalization constant to *c* = *ĉ* = max_*y*,*θ*_ *π*(*D*|*y*, *θ*) where it is ignored whether the model is able to simulate such values, as in the initial approach by Wilkinson (2013). In addition, we compared ASSA to the noisy model sampler A2 in a sequential form. We stopped runs when the acceptance rate fell below 1e-3, or exact inference with *T* = 1 was achieved.

For the deterministic models M1 and M2, we could confirm that the distributions inferred by ASSA closely matched the theoretical ones (Supplementary Information, Figure S9). Unfortunately, such an analysis is not that easily possible for the stochastic models M3 and M4. Yet, the posterior distributions obtained using ASSA for all four models are centered around the true parameters to a degree that seems reasonable given model and data (Figure 4A). When other samplers of type A3 reached *T* = 1, the estimated posteriors were similar in shape. The posterior approximations obtained by the noisy model sampler A2 differed for three models from those obtained by ASSA, indicating that this approach is not able to yield reliable results within a reasonable computational budget.

**Figure 4:**
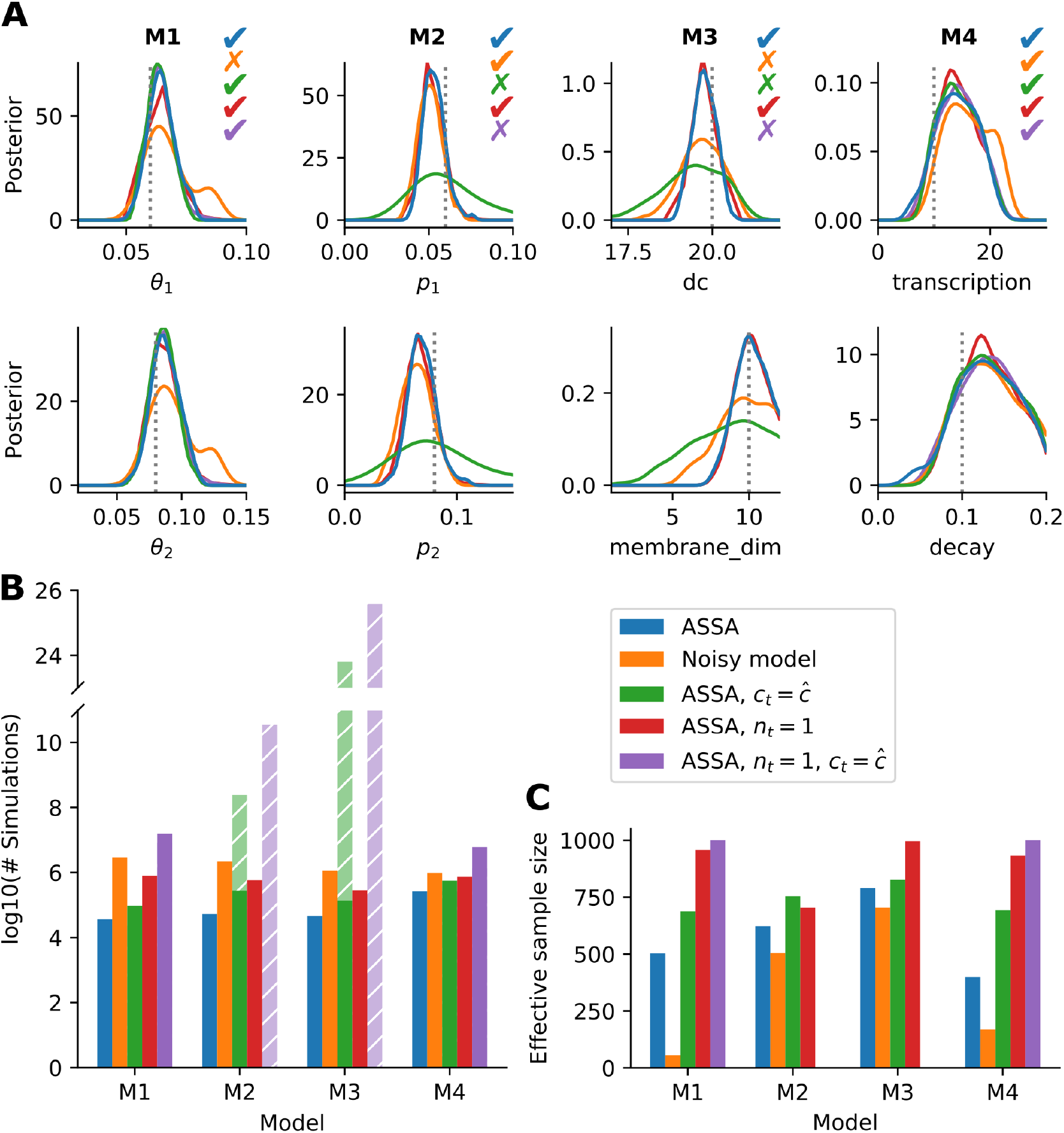
Method comparison for models M1-4. (A) Posterior marginals obtained using the five different samplers. For runs that had to be stopped due to an acceptance rate below 1e-3, the last finished population is shown. True parameters are indicated by dotted lines. The colored check marks indicate the methods reaching *T* = 1, or for the noisy model an adequate approximation. (B) Total number of simulations over all iterations. For samplers that had to be stopped early, in addition the estimated values are shown hatched. (C) Effective sample sizes for the final posterior estimate obtained by each algorithm in its last iteration.

### 3.6 Substantial speedup compared to established approaches

A commonly used measure for efficiency of a sampler is the total number of required model simulations. This is for dynamical models typically the time-critical part (see e.g. Jagiella *et al*. (2017)). We found that for all models, ASSA required the least simulations (Figure 4B). For model M4 the advantage was with a factor of 2 the smallest, which could be explained by the model itself being very stochastic, so that simulating the noisy data was not unlikely. The other models possessed more internal structure, i.e. a higher signal-to-noise ratio, such that model simulations close to the noisy data were rather unlikely. For models M2 and M3, ASSA with *c* = *ĉ*, as well as the version with *n_t_* = 1, was not even able to reach *T* = 1 in the computational budget, resulting in a higher posterior variance due to an overestimation of the noise variance (see Figure 4A).

For our analysis, we provided all samplers with a computational budget which was by far sufficient for ASSA. Unfortunately, for some runs with *c* = *ĉ*, this budget was still insufficient. To compare the expected computational budget required for exact sampling, we made use of the roughly inverse proportional dependence of the acceptance rate on the normalization factor *c* (3). With s the number of simulations required by ASSA in the last iteration, we get the estimate 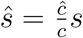 for the required number of simulations for samplers using *c* = *ĉ* in the last iteration. The estimates indicate that the sequential samplers with *c* = *ĉ* would require 4 and 20 orders of magnitude more simulations than those with self-tuned *c*, on model M2 and M3, respectively (Figure 4B). Even with massive parallelization, exact inference would thus not be possible without the proposed self-tuning scheme for *c*.

On all models M1-4, ASSA required about 22, 11, 6, and 2 times less simulations, respectively, than with *n_t_* = 1, indicating that the temperature selection scheme allows to efficiently bridge from prior to posterior.

To assess the influence of the number of data points, we performed the inference for models M1-3 for 10 to 1000 data points (Supplementary Information, Section 8.3). The value of *ĉ* grew exponentially with the number of data points, while this was not the case for the highest *c* possible under the model, 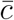. Indeed, we found for models M1-3 that for ASSA the number of simulations increased only moderately. The ratio *ĉ*/*c* with *c* the value used by ASSA in the last iteration increased e.g. for model M1 from about 10 for 10 data points to more than 1e200 for 1000 data points. This indicates that ASSA scales well with the number of data points, while approaches with too large *c* quickly become computationally infeasible.

While the number of required model evaluations provides information about the complexity, it does not account for stability. This is assessed with the effective sample size 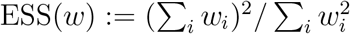 (Martino *et al*., 2017), with *w_i_* denoting the importance weights. The ESS is a heuristic measure of how many independent samples a population of particles obtained via importance sampling effectively consists of, decreasing for more volatile weights. Since we inflate the weights by (2) if the criterion exceeds 1, it is to be expected that ASSA has a lower ESS than the alternative stochastic acceptors, which indeed was the case on the test models (Figure 4C). However, for all models the ESS was still reasonably high, indicating that the population is not degenerated. E.g. on model M3 an increase of the population size by 25% would presumably yield an ESS of more than 1000. Compared to the orders of magnitude differences between sample numbers, this renders ASSA highly efficient (Supplementary Information, Figure S13).

### 3.7 Scales to challenging estimation problems

To assess the performance of the proposed approach in practice, we considered the application problems M5 and M6. Since the acceptance rates when updating *c* always to the highest so far observed value were too low, once acceptance rates fell below 0.1 we employed a factor *β* of 20 and 10, respectively, as defined in Section 2.6.

For model M5, we used a population size of *N* = 1*e*4 to guarantee stability of the results. The posterior marginals (Figure 5) indicate that seven parameters can be accurately estimated, two parameters can be constrained, and two parameters are non-identifiable. The posterior distributions recover the reported literature values (Boehm *et al*., 2014), which had been obtained by optimization, and the parameter samples provide an accurate description of the experimental data (Supplementary Information, Figure S14). Importantly, in addition to kinetic parameters our approach was able to identify the standard deviations of the normal noise models on all three parameters. In contrast, the application of a sequential version of the noisy model approach A2 performed worse and was in particular unable to fit all noise parameters. The ESS for the stochastic acceptor was 1011 (using in total 8e6 simulations), for the noisy model only 83 (using in total 12e6 simulations).

**Figure 5:**
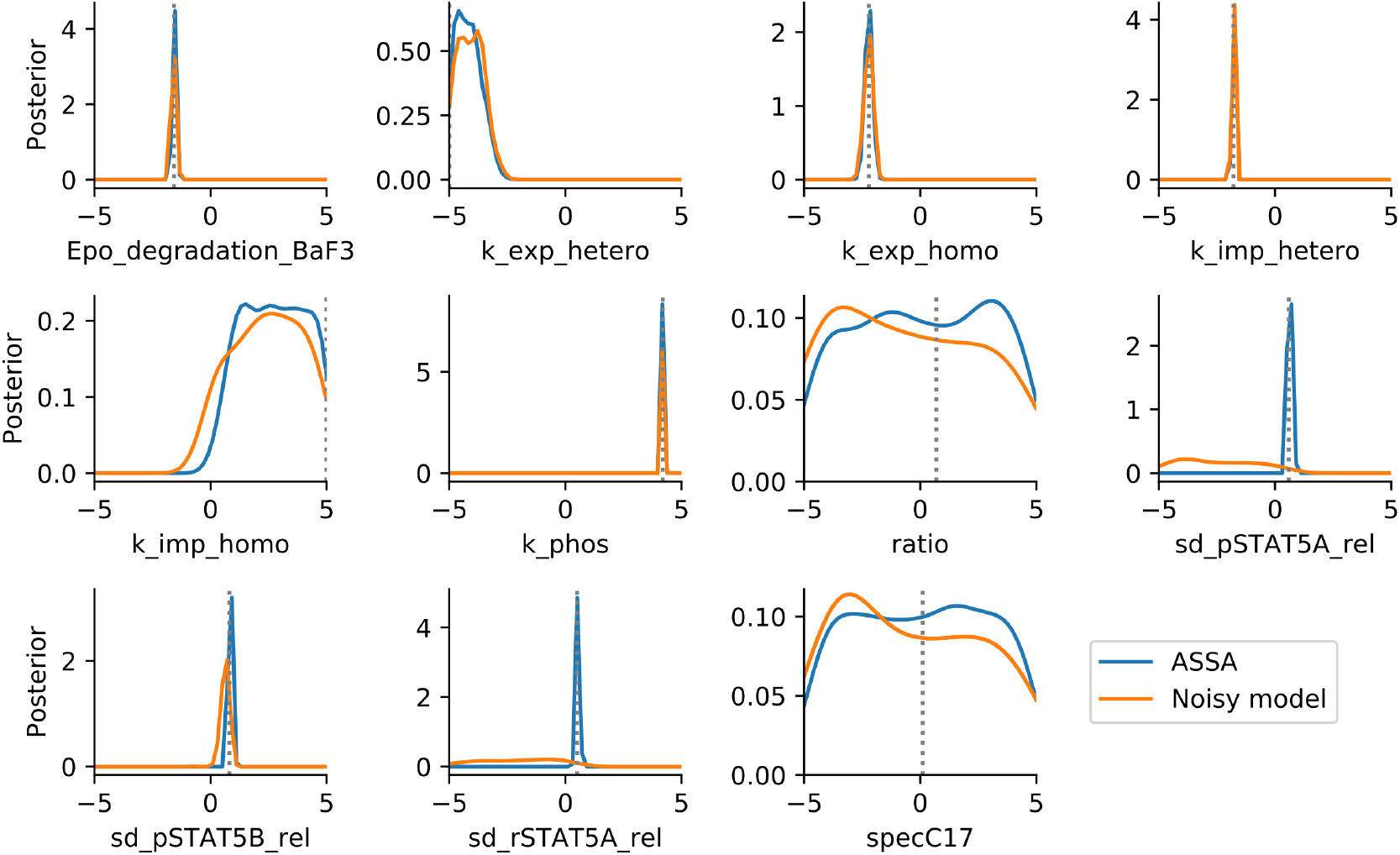
Posterior marginals for model M5, using ASSA and the noisy model sampler. The literature MAP values are indicated by dotted lines.

For model M6, we used a population size of *N* = 5*e*2 due to the high simulation time of the model. Again, the comparison of the posterior marginals with the noisy model, which was given a similar computational budget as required by the stochastic acceptor, reveals that the stochastic acceptor extracted overall more information (Supplementary Information, Figure S15), and the parameter samples describe the considered data accurately under the assumed noise model (Supplementary Information, Figure 16). Given the data, the parameters related to initial conditions cannot be inferred well, but for the others the reference values are accurately matched. The numbers of simulations were 4.0e5 and 3.2e5, with an ESS of 222 and 153, respectively for ASSA and the noisy model sampler.

## 4 Discussion

Modern ABC and ABC-SMC algorithms allow to perform parameter estimation for complex models. However, while easy to apply, these algorithms can lead to wrong results if measurement noise is not correctly accounted for. Here, we discussed ways of dealing with noise and presented an adaptive sequential importance sampling algorithm (ASSA), broadly applicable to various models and noise models. We demonstrated, using several test models, that the proposed algorithm is more accurate and up to orders of magnitude more efficient than existing approaches. We achieved this efficiency gain by learning a required normalization constant in a self-tuned manner, correcting for potential biases by reweighting, and by devising an adaptive tempering scheme which in particular allows to find a good starting temperature. Thence, we were able to perform exact likelihood-free inference on models on which this would have hitherto been impossible. Further, we implemented the algorithm in the freely available toolbox pyABC.

Our approach is self-adaptive to the problem structure by learning good values for several tuning parameters such as the initial temperature and the normalization constant. By learning proper values for these parameters on the fly, our approach is both stable and applicable to diverse problems, and gets rid of the need of manual tuning, which otherwise can be time-intensive. This is a key difference to related approaches (e.g. Daly *et al*. (2017)), in which parameters are manually adapted to the individual problems. Furthermore, the proposed approaches allows to estimate noise levels, facilitating integrated analysis workflows (Hross *et al*., 2018).

Given that ABC has in the past been frequently applied without a proper noise formulation, the question may be raised whether this may have lead to wrong parameter estimates. Unfortunately, this is difficult to answer. However, since we have been able to apply exact likelihood-free inference even to computationally demanding problems, we expect the algorithm to be broadly applicable in the future, thus improving the reliability of parameter estimates for a broad spectrum of applications.

One possible path of future research is to investigate improved tempering schemes. The overall required number of simulations depends both on the number of intermediate populations, and the acceptance rates between steps. The exponential decay scheme we employed was rather loosely motivated by analogies to parallel tempering MCMC. There exist approaches to dynamically adjust the temperature steps (Predescu *et al*., 2004; Vousden *et al*., 2015), however these are specific to parallel tempering MCMC. For likelihood based SMC, there exist approaches that try to keep the effective sample size constant (Latz *et al*., 2018), however it remains to be investigated whether these are applicable in a likelihood-free context. Another interesting use of tempering is for thermodynamic integration, allowing to compute Bayes factors. Thus, the presented algorithm could potentially be extended to allow to perform model selection, as an alternative to e.g. Toni and Stumpf (2010).

In conclusion, our results demonstrate the importance and the benefits of using proper noise models. The proposed algorithms can exploit the structure of the noise model to perform exact inference for computationally demanding models. As the implementation in pyABC facilitates massive parallelization, this approach is also applicable to computationally demanding problems.

## Author contributions

Both authors derived the theoretical foundation and devised the algorithms. Y.S. wrote the implementation and performed the case study. Both authors discussed the results and conclusions and jointly wrote and approved the final manuscript.

## References

Andrieu, C., Doucet, A., and Holenstein, R. (2010). Particle markov chain monte carlo methods. Journal of the Royal Statistical Society: Series B (Statistical Methodology), 72(3), 269–342.

Banga, J. R. (2008). Optimization in computational systems biology. BMC Syst. Biol., 2(47).

Beaumont, M. A., Zhang, W., and Balding, D. J. (2002). Approximate Bayesian Computation in population genetics. Genetics, 162(4), 2025–2035.

Beaumont, M. A., Cornuet, J.-M., Marin, J.-M., and Robert, C. P. (2009). Adaptive approximate bayesian computation. Biometrika, 96(4), 983–990.

Boehm, M. E., Adlung, L., Schilling, M., Roth, S., Klingmueller, U., and Lehmann, W. D. (2014). Identification of isoform-specific dynamics in phosphorylation-dependent stat5 dimerization by quantitative mass spectrometry and mathematical modeling. Journal of proteome research, 13(12), 5685–5694.

Coxe, S., West, S. G., and Aiken, L. S. (2009). The analysis of count data: A gentle introduction to poisson regression and its alternatives. Journal of personality assessment, 91(2), 121–136.

Daly, A. C., Cooper, J., Gavaghan, D. J., and Holmes, C. (2017). Comparing two sequential monte carlo samplers for exact and approximate bayesian inference on biological models. Journal of The Royal Society Interface, 14(134), 20170340.

Del Moral, P., Doucet, A., and Jasra, A. (2006). Sequential monte carlo samplers. Journal of the Royal Statistical Society: Series B (Statistical Methodology), 68(3), 411–436.

Del Moral, P., Doucet, A., and Jasra, A. (2012). An adaptive sequential monte carlo method for approximate bayesian computation. Statistics and Computing, 22(5), 1009–1020.

Drovandi, C. C. (2018). Abc and indirect inference. In Handbook of Approximate Bayesian Computation. CRC Press (Taylor & Francis Group).

Earl, D. J. and Deem, M. W. (2005). Parallel tempering: Theory, applications, and new perspectives. Physical Chemistry Chemical Physics, 7(23), 3910–3916.

Eriksson, O., Jauhiainen, A., Maad Sasane, S., Kramer, A., Nair, A. G., Sartorius, C., and Hellgren Kotaleski, J. (2019). Uncertainty quantification, propagation and characterization by bayesian analysis combined with global sensitivity analysis applied to dynamical intracellular pathway models. Bioinformatics, 35(2), 284–292.

Fearnhead, P. and Prangle, D. (2012). Constructing summary statistics for approximate bayesian computation: semi-automatic approximate bayesian computation. Journal of the Royal Statistical Society: Series B (Statistical Methodology), 74(3), 419–474.

Frazier, D. T., Robert, C. P., and Rousseau, J. (2020). Model misspecification in approximate bayesian computation: consequences and diagnostics. Journal of the Royal Statistical Society: Series B (Statistical Methodology).

Gillespie, D. T. (1977). Exact stochastic simulation of coupled chemical reactions. The journal of physical chemistry, 81(25), 2340–2361.

Goldwyn, J. H., Imennov, N. S., Famulare, M., and Shea-Brown, E. (2011). Stochastic differential equation models for ion channel noise in hodgkin-huxley neurons. Physical Review E, 83(4), 041908.

Gourieroux, C., Monfort, A., and Renault, E. (1993). Indirect inference. Journal of applied econometrics, 8(S1), S85–S118.

Hines, K. (2015). A primer on bayesian inference for biophysical systems. Biophysical Journal, 108(9), 2103–2113.

Hross, S., Theis, F. J., Sixt, M., and Hasenauer, J. (2018). Mechanistic description of spatial processes using integrative modelling of noise-corrupted imaging data. J. R. Soc. Interface, 15(149), 20180600.

Imle, A., Kumberger, P., Schnellbächer, N. D., Fehr, J., Carrillo-Bustamante, P., Ales, J., Schmidt, P., Ritter, C., Godinez, W. J., Müller, B., et al. (2019). Experimental and computational analyses reveal that environmental restrictions shape hiv-1 spread in 3d cultures. Nature communications, 10(1), 2144.

Jagiella, N., Rickert, D., Theis, F. J., and Hasenauer, J. (2017). Parallelization and high-performance computing enables automated statistical inference of multi-scale models. Cell Systems, 4(2), 194–206.

Klinger, E. and Hasenauer, J. (2017). A scheme for adaptive selection of population sizes in Approximate Bayesian computation - Sequential Monte Carlo. In J. Feret and H. Koeppl, editors, Computational Methods in Systems Biology. CMSB 2017, volume 10545 of Lecture Notes in Computer Science. Springer, Cham.

Klinger, E., Rickert, D., and Hasenauer, J. (2018). pyABC: distributed, likelihood-free inference. Bioinformatics, 34(20), 3591–3593.

Latz, J., Papaioannou, I., and Ullmann, E. (2018). Multilevel sequential2 monte carlo for bayesian inverse problems. Journal of Computational Physics, 368, 154–178.

Lenive, O., Kirk, P. D., and Stumpf, M. P. (2016). Inferring extrinsic noise from single-cell gene expression data using approximate bayesian computation. BMC systems biology, 10(1), 81.

Liu, J. S., Chen, R., and Wong, W. H. (1998). Rejection control and sequential importance sampling. Journal of the American Statistical Association, 93(443), 1022–1031.

Maier, C., Loos, C., and Hasenauer, J. (2017). Robust parameter estimation for dynamical systems from outlier-corrupted data. Bioinformatics, 33(5), 718–725.

Martino, L., Elvira, V., and Louzada, F. (2017). Effective sample size for importance sampling based on discrepancy measures. Signal Processing, 131, 386–401.

Picchini, U. (2014). Inference for sde models via approximate bayesian computation. Journal of Computational and Graphical Statistics, 23(4), 1080–1100.

Prangle, D. et al. (2017). Adapting the abc distance function. Bayesian Analysis, 12(1), 289–309.

Predescu, C., Predescu, M., and Ciobanu, C. V. (2004). The incomplete beta function law for parallel tempering sampling of classical canonical systems. The Journal of chemical physics, 120(9), 4119–4128.

Price, L. F., Drovandi, C. C., Lee, A., and Nott, D. J. (2018). Bayesian synthetic likelihood. Journal of Computational and Graphical Statistics, 27(1), 1–11.

Raue, A., Kreutz, C., Maiwald, T., Bachmann, J., Schilling, M., Klingmüller, U., and Timmer, J. (2009). Structural and practical identifiability analysis of partially observed dynamical models by exploiting the profile likelihood. Bioinformatics, 25(25), 1923–1929.

Raue, A., Schilling, M., Bachmann, J., Matteson, A., Schelke, M., Kaschek, D., Hug, S., Kreutz, C., Harms, B. D., Theis, F. J., Klingmüller, U., and Timmer, J. (2013). Lessons learned from quantitative dynamical modeling in systems biology. PLoS ONE, 8(9), e74335.

Sisson, S. and Fan, Y. (2018). Abc samplers. In Handbook of Approximate Bayesian Computation, pages 87–123. Chapman and Hall/CRC.

Sisson, S., Fan, Y., and Beaumont, M. (2018a). Overview of abc. Handbook of Approximate Bayesian Computation, pages 3–54.

Sisson, S. A., Fan, Y., and Tanaka, M. M. (2007). Sequential monte carlo without likelihoods. Proceedings of the National Academy of Sciences, 104(6), 1760–1765.

Sisson, S. A., Fan, Y., and Beaumont, M. (2018b). Handbook of approximate Bayesian computation. Chapman and Hall/CRC.

Sugita, Y., Kitao, A., and Okamoto, Y. (2000). Multidimensional replica-exchange method for free-energy calculations. The Journal of Chemical Physics, 113(15), 6042–6051.

Toni, T. and Stumpf, M. P. H. (2010). Simulation-based model selection for dynamical systems in systems and population biology. Bioinformatics, 26(1), 104–110.

Toni, T., Welch, D., Strelkowa, N., Ipsen, A., and Stumpf, M. P. H. (2009). Approximate Bayesian computation scheme for parameter inference and model selection in dynamical systems. J. R. Soc. Interface, 6, 187–202.

van der Vaart, E., Prangle, D., and Sibly, R. M. (2018). Taking error into account when fitting models using approximate bayesian computation. Ecological applications, 28(2), 267–274.

Vousden, W., Farr, W. M., and Mandel, I. (2015). Dynamic temperature selection for parallel tempering in markov chain monte carlo simulations. Monthly Notices of the Royal Astronomical Society, 455(2), 1919–1937.

Wilkinson, R. D. (2013). Approximate bayesian computation (abc) gives exact results under the assumption of model error. Statistical applications in genetics and molecular biology, 12(2), 129–141.

Wood, S. N. (2010). Statistical inference for noisy nonlinear ecological dynamic systems. Nature, 466(7310), 1102.

