## Supplementary Information for "Efficient Exact Inference for Dynamical Systems with Noisy Measurements using Sequential Approximate Bayesian Computation"

January 2020

### Overview

This supplementary information is structured as follows: In Chapter 1, we shortly formulate the ABC problem and also discuss the extension to general acceptance kernels. In Chapter 2, we discuss the problem of measurement noise in ABC and show how one can in theory correct for it. In Chapter 3, we then present details on our proposed adaptive sequential approach. Chapter 4 contains details on how certain normalization constants were computed. Chapter 5 describes the implementation. In Chapter 6, we give examples of wrong analyses in the presence of noise. In Chapter 7, we give details on the test models used in the main manuscript, and Chapter 8 contains additional analyses and figures supplementing the discussion on the test models in the main manuscript.

### 1 Approximate Bayesian Computation

#### 1.1 The problem

Throughout this document, we consider parameters $\theta \in \mathbb{R}^{n_\theta}$, a (Lebesgue) prior density $\pi(\theta)$ encoding prior information, and observed data $D \in \mathbb{R}^{n_y}$. Further, we consider a model likelihood $p(y|\theta)$, the density of the model having output data $y$ given model parameterization $\theta$. Together, according to Bayes' theorem these yield the posterior density of parameters given the observed data,

$$\pi(\theta|D) = \frac{p(D|\theta)\pi(\theta)}{\pi(D)} \propto p(D|\theta)\pi(\theta),$$

where the evidence $\pi(D) := \int p(D|\theta)\pi(\theta)\, d\theta$ is just a normalization constant that can be ignored in common Monte Carlo techniques.

In the first part, we assume that $D \sim p(y|\theta_{\text{true}})$, i.e. that for some unknown parameterization $\theta_{\text{true}}$, the model correctly describes the data generation process, i.e. $D$ was generated as an instance of the model. When we discuss the influence of model error, we instead assume that $D \sim q(\bar{y}|\theta_{\text{true}})$ has been generated under a different model.

#### 1.2 Standard Rejection ABC

We assume that while we can sample from the likelihood function simulated data $y \sim p(y|\theta)$, we cannot evaluate the likelihood itself. The idea of Approximate Bayesian Computation (ABC) is to replace this by assessing the similarity of simulated and observed data. Therefore, in its most common form, *Rejection ABC*, one defines a distance metric $d : \mathbb{R}^{n_y} \times \mathbb{R}^{n_y} \to \mathbb{R}$ and a threshold $\varepsilon > 0$, and proceeds as follows:

1. sample parameters $\theta \sim \pi(\theta)$,
2. simulate data $y \sim p(y|\theta)$,
3. accept if $d(y, D) \leq \varepsilon$.

This is repeated until sufficiently many particles $\{\theta_i\}_{i=1,\dots,N}$ have been accepted. This way, the distribution we are sampling from as an approximation of the true posterior distribution is

$$\pi_{\text{ABC},\varepsilon}(\theta|D) \propto \int I(d(y, D) \leq \varepsilon) p(y|\theta)\, dy \cdot \pi(\theta), \tag{1}$$

where $I$ denotes the indicator function. Note that due to the course of dimensionality, in ABC commonly summary statistics $s = s(y)$ are employed. Here, we abstract from this by assuming $s = y$ has already been incorporated if applicable. This suffices for our purposes, but raises some questions, in particular on whether later we assume a noise model on the data, or the summary statistics.

#### 1.3 General acceptance kernels

This algorithm can be generalized by replacing the acceptance step by a generic acceptance kernel, i.e. a non-negative integrable function $K_\varepsilon : \mathbb{R}^{n_y} \to \mathbb{R}$, $y \mapsto K_\varepsilon(D|y)$, s.t. $\int K(D|y)\, dy = 1$, and defining

$$\pi_{\text{ABC},\varepsilon}(\theta|D) \propto \int K_\varepsilon(D|y) p(y|\theta)\, dy \cdot \pi(\theta). \tag{2}$$

This distribution is sampled from by replacing the third step above by

3. accept with probability $\frac{K_\varepsilon(y)}{\sup_{y^*} K_\varepsilon(y^*)}$.

$K_\varepsilon$ can in particular be chosen to give a higher weight to simulations that are close to the observed data, and usually gets more centered around $D$ for $\varepsilon \to 0$. Note that with $K_\varepsilon(D|y) \propto I(d(y, D) \leq \varepsilon)$ (the normalization does not matter to us in sampling), the indicator function can be included in this formulation, and is also referred to as *uniform kernel*.

Commonly, a series of kernels for arbitrary $\varepsilon > 0$ is constructed based on a non-negative integrable function $K : \mathbb{R}^{n_y} \to \mathbb{R}$, $y \mapsto K(y)$, s.t. $\int K(y)\, dy = 1$, and then defining $K_\varepsilon(y) := \frac{1}{\varepsilon^{n_y}} K(\frac{y-D}{\varepsilon})$.

#### 1.4 ABC gives asymptotically exact inference

ABC introduces an approximation error by accepting simulated data that do not exactly match the observed data. In the following we verify (see e.g. also the derivations in Sisson et al. [2018]), that if we let $\varepsilon \to 0$, in the limit only accepting if the simulated data $y$ exactly match the observed data $D$, we have inference from the true posterior distribution, i.e.

$$\pi_{\text{ABC}}(\theta|D) := \lim_{\varepsilon \to 0} \pi_{\text{ABC},\varepsilon}(\theta|D) = \pi(\theta|D).$$

We require that the acceptance kernels form a Dirac delta sequence.

**Theorem 1.** *Define the ABC posterior distribution as in* (2)*, assume that* $D \sim p(y|\theta)$*, and that* $\{K_\varepsilon\}_{\varepsilon \to 0}$ *is a Dirac delta sequence centered at* $D$*. Then,*

$$\pi_{\text{ABC},\varepsilon}(\theta|D) \to \pi(\theta|D), \quad \varepsilon \to 0. \tag{3}$$

*Proof.* It is

$$\lim_{\varepsilon \to 0} \pi_{\text{ABC},\varepsilon}(\theta|D) \propto \lim_{\varepsilon \to 0} \int K_\varepsilon(D|y) p(y|\theta)\, dy \cdot \pi(\theta) = \int \delta(y = D) p(y|\theta)\, dy \cdot \pi(\theta) = p(D|\theta)\pi(\theta).$$

□

A core assumption here is that $D \sim p(y|\theta)$, i.e. that $p$ captures the entire data generation process. Otherwise, as we see in the subsequent section, we have convergence to a wrong posterior.

For the kernels commonly used in ABC, it is straightforward to show that these indeed give rise to Dirac delta sequences: For the kernel-induced sequence note the following

**Lemma 1** (Generating functions of Dirac delta sequences)**.** *Let* $K : \mathbb{R}^n \to \mathbb{R}$ *be non-negative, Lebesgue integrable with* $\int K(x)\, dx = 1$*. Then the sequence of functions* $K_\varepsilon(x) := \frac{1}{\varepsilon^{n_y}} K(\frac{x}{\varepsilon})$ *converges to the Dirac delta distribution as* $\varepsilon \to 0$ *in the sense of generalized functions.*

*Proof.* See Kanwal [2012]. □

For the distance based form (1), the statement trivially follows as above if the distance measure is norm induced, $d(y, D) = \|y - D\|^p$. The general setting is discussed in Prangle [2017] based on the Lebesgue differentiation theorem.

#### 1.5 Model error in ABC

In practice, it is in general not possible to have $\varepsilon \to 0$, at least in the case of continuous data and a non-deterministic model, as samples must exactly replicate the observed data, the probability for which is in general 0. For a given $\varepsilon \geq 0$ and kernel $K_\varepsilon$, in ABC we effectively generate samples from a model assuming that the observed data $D$ are distributed as

$$\pi_{\text{ABC},\varepsilon}(D|\theta) \propto \int K_\varepsilon(D|y)p(y|\theta)\,dy\,.$$

In the most common application of a norm-induced distance $d(y, D) = \|y - D\|$ and using a uniform kernel, this becomes

$$\pi_{\text{ABC},\varepsilon}(D|\theta) \propto \int I(\|y - D\| \leq \varepsilon)p(y|\theta)\,dy,$$

i.e. we effectively assume the data to be a convolution of the original model with the acceptance kernel.

Now let us assume that the model does not accurately describe the data generation process, i.e. we assume a model $D \sim p(y|\theta)$ but in real $D \sim q(\bar{y}|\theta)$. Then obviously we perform analysis for the wrong model. The same problem exists in likelihood-based inference. However, this problem is more severe in ABC, because ABC is based on simulating data and comparing them via some distance measure to the observed data. In particular it is thus easy and convenient to completely ignore measurement noise, only focusing on the mechanistic, noise-free, data generation process. This would not be possible in likelihood-based inference, which is commonly based on a non-degenerate noise model in order to formulate the likelihood.

A particular problem occurs when the used model $p$ is not, or only with a small probability, able to explain the observed data. In practice, this is easily possible if the model does not account for measurement noise, because a mechanistic model usually possesses some internal structure, while e.g. noise tends to be uncorrelated among time points. In likelihood-based inference, $p(D|\theta) = 0$ would immediately reveal that the model inadequacy. However, in ABC this information is not available. Assuming the above uniform kernel, $p(D|\theta) = 0$ will only become apparent through $\varepsilon$ being unable to fall below some $\varepsilon_{\min} > 0$, such that the distribution for $D$ converges to

$$\pi_{\text{ABC},\varepsilon_{\min}}(D|\theta) \propto \int I(\|y - D\| \leq \varepsilon_{\min})p(y|\theta)\,dy\,.$$

In practice, this form of model error is not easy to detect, because even if the model is correct, computational resources usually require the analysis to be stopped at some $\varepsilon$.

It has been argued that the exact distance metric is of minor importance in ABC [McKinley et al., 2009, Owen et al., 2015], except for efficiency improvements [Prangle, 2017]. However, that only holds for $\varepsilon \to 0$. If $\varepsilon_{\min} \gg 0$, the distance metric does matter, as it defines the domain of simulations yielding similar distances. In particular, for a deterministic model, i.e. $p(y|\theta) = \delta(y - y(\theta))$, we find

$$\pi_{\text{ABC},\varepsilon_{\min}}(\theta|y_{\text{obs}}) \propto I(d(y(\theta), y_{\text{obs}}) = \varepsilon_{\min}) \cdot \pi(\theta).$$

This can be interpreted as a distribution over all parameters $\theta$ whose simulated data reproduce the overall minimum distance to the observed data, weighted by the prior. What ABC effectively does in this situation is minimize the distance function, and only points that have that minimum distance are accepted. Now, for example, a weighted $\ell_2$ distance

$$d(y, D) = \sum_i \omega_i(y_i - D_i)^2)$$

can be interpreted as the negative log-likelihood of a normal distribution

$$p(D|y, \theta) \propto \bigotimes_i \mathcal{N}(D_i|y_i, \omega_i^{-1}),$$

but we do *not* obtain the full Bayesian posterior assuming this noise model, but only those points giving the *maximum likelihood value* under the assumption of that normal distribution, weighted by the prior. If we instead use e.g. an $\ell_1$ distance

$$d(y, D) = \sum_i \omega_i |y_i - D_i|,$$

we effectively obtain maximum likelihood points assuming a Laplace noise model

$$p(D|y, \theta) \propto \bigotimes_i \text{Laplace}(D_i | y_i, \omega_i^{-1}).$$

So, while some distances effectively implicitly translate to some noise distribution, parameters giving the maximum likelihood value and weighted by the prior is far from what we aspire in a Bayesian analysis. To obtain maximum likelihood estimates, there exist far more efficient approaches

#### 2 Exact inference in ABC assuming measurement noise

In the following, we assume the observed data to be noisy. That is, we assume that the data are not a realization of $p(y|\theta_{\text{true}})$, but of $q(\bar{y}|\theta_{\text{true}})$. In that case, assuming likelihood $p$ in Section 1 simply implies that we do inference for the wrong model. Corresponding to the notion of noise, we further assume that we can write $(\bar{y}, y) \sim \pi(\bar{y}|y,\theta)p(y|\theta)$, so that $q(\bar{y}|\theta) \propto \int \pi(\bar{y}|y,\theta)p(y|\theta)\,dy$. The interpretation of $\pi(\bar{y}|y,\theta)$ is that of a parameterized noise model of observing data $\bar{y}$ under noise-free model output $y$ and parameters $\theta$. Thus, the noise model relates observables that assume perfect measurements to practically obtained noisy data. We call $\pi(\bar{y}|y,\theta)$ the *noise model*, $p(y|\theta)$ the *model likelihood* and $q(\bar{y}|\theta)$ the *full likelihood*.

The goal is now to infer the corrected posterior

$$\pi(\theta|D) \propto q(D|\theta)\pi(\theta) \propto \int \pi(D|y,\theta)p(y|\theta)\,dy \cdot \pi(\theta).$$

We assume that we can evaluate the noise model, while we may only be able to sample from but not evaluate the model likelihood. The idea is to keep the first two steps, i.e. to sample parameters $\theta \sim \pi(\theta)$ and noise-free data $y \sim p(y|\theta)$, but to replace the acceptance step by

3. accept with probability $\frac{\pi(D|y,\theta)}{c}$ where $c \geq \sup_{y,\theta} \pi(D|y,\theta)$.

This step can be implemented as follows: Sample $u \sim U[0,1]$ and accept if $\frac{\pi(D|y,\theta)}{c} \geq u$.

The following theorem tells that the noise model allows to do exact inference even though not being able to evaluate the model likelihood. The algorithm

**Theorem 2** (Exact noisy ABC)**.** *Consider a prior density $\pi(\theta)$, a likelihood of noise-free model outputs given parameters $p(y|\theta)$, and a likelihood of data given model outputs and parameters $\pi(\bar{y}|y,\theta)$, and assume $D \sim q(\bar{y}|\theta) \propto \int \pi(\bar{y}|y,\theta)p(y|\theta)\,dy$. Then the above procedure with $c \geq \hat{c} := \sup_{y,\theta} \pi(D|y,\theta)$ targets the correct posterior distribution $\pi(\theta|D) \propto q(D|\theta)\pi(\theta)$.*

*Proof.* This proof is adapted from Wilkinson [2013]. Let

$$A = \begin{cases} 1 & \text{if } \theta \text{ is accepted,} \\ 0 & \text{otherwise.} \end{cases}$$

Then for arbitrary $c > 0$ the distribution of accepted parameters is given by

$$\pi(\theta|A=1) = \frac{\pi(A=1|\theta)\cdot\pi(\theta)}{\pi(A=1)} = \frac{\int \min\left[\frac{\pi(D|y,\theta)}{c}, 1\right] p(y|\theta)\,dy \cdot \pi(\theta)}{\iint \min\left[\frac{\pi(D|y,\theta')}{c}, 1\right] p(y|\theta')\,dy \cdot \pi(\theta')\,d\theta'},$$

using that

$$\pi(A=1|\theta) = \int \min\left[\frac{\pi(D|y,\theta)}{c}, 1\right] p(y|\theta)\,dy\,.$$

Moreover, since we assume the data to be generated under $D \sim q(\bar{y}|\theta)$, the true posterior distribution is given by

$$\pi(\theta|D) = \frac{q(D|\theta)\pi(\theta)}{\pi(D)} = \frac{\int \pi(D|y,\theta)p(y|\theta)\,dy \cdot \pi(\theta)}{\iint \pi(D|y,\theta')\pi(y|\theta')\,dy \cdot \pi(\theta')\,d\theta'},$$

using that

$$q(D|\theta) \propto \int \pi(D|y,\theta)p(y|\theta)\,dy\,.$$

Thus, if $c \geq \sup_{y,\theta} \pi(D|y,\theta)$, we find $\pi(\theta|A=1) = \pi(D|\theta)$, i.e. the distribution we sample from is the true posterior distribution. □

*Remark* 1. Our formulation allows conditioning the noise model $\pi(\bar{y}|y,\theta)$ on the parameter $\theta$, yielding a parameterized noise model. This is useful if the noise distribution depends on parameters that are not known. An example are variance parameters, which are often not known in practice [Raue et al., 2013]. Including these as parameters in the parameter estimation problem allows to infer them alongside the dynamical model parameters. An intuitive explanation for why e.g. variance parameters are not estimated arbitrarily large is that, while allowing to accept simulations that do not tightly match the observed data, large variances are implicitly punished by a flatter density, leading to smaller acceptance rates.

*Remark* 2. If $c$ is too small s.t. higher values occur during sampling, the effective noise distribution sampled from is not $\pi(D|y,\theta)$, but $\min\left[\frac{\pi(D|y,\theta)}{c}, 1\right]$, i.e. it is flattened out to a uniform distribution at values of high probability. In particular as $c \searrow 0$, the effective noise distribution becomes one similar to a uniform distribution of increasing variance. For a more illustrative discussion of this, see Daly et al. [2017]. By reweighting, the algorithm we propose corrects for this flattening.

*Remark* 3. It suffices if

$$c \geq \bar{c} := \sup\left\{\pi(D|y,\theta) \mid p(y|\theta) > 0, \pi(\theta) > 0)\right\},$$

that is the noise models for all $(y,\theta)$ that are actually possible under the model. In fact, it is even sufficient if $c$ is higher than all $\pi(D|y,\theta)$ generated in the sampling process. This has practical implementations, because usually the highest peak of the noise model is at $\hat{c} = \pi(D|D,\theta)$ for some variance minimal $\theta$, i.e. when the simulated data match the observed data. However, it will in general not be possible or only with a small probability to generate data $y \approx D$ under the noise-free model likelihood. This has practical implications, because it allows $c$ to be smaller, yielding higher acceptance rates. A problem is that while $\hat{c}$ is generally easy to compute, $\bar{c}$ is in general intractable. This is why we propose a method to estimate $\bar{c}$ automatically in a Sequential Monte Carlo algorithm.

*Remark* 4. Theorem 2 can also be seen in light of Section 1.3, using the noise model as a generalized acceptance kernel. However, the distinction is that with the noise model we are not interested in reducing the kernel variance to 0, which is necessary in Section 1.3 to have asymptotically exact inference. It can be stated that the non-degenerate noise allows us to do exact likelihood-free inference.

*Remark* 5. Since $\int \pi(D|y,\theta)p(y|\theta)\,dy$ defines a probability density, in particular $\pi(D|y,\theta)p(y|\theta)$ is integrable and thus defines a probability density for $y$, which we use implicitly.

### 3 Towards an efficient exact sequential ABC sampler

#### 3.1 ABC-SMC

ABC is commonly integrated with a sequential Monte Carlo (ABC-SMC) scheme to increase efficiency. In ABC-SMC, the Rejection ABC steps are iterated over a number of generations $t = 1,\dots,n_t$. In generation $t$, parameters $\theta \sim g_t(\theta)$ are sampled from some proposal distribution $g_t(\theta) \gg \pi(\theta)$, usually based on the accepted particles of the previous generation, with $g_1(\theta) = \pi(\theta)$, s.t. over time the proposal distribution more closely resembles the posterior distribution. Then, for simulated data $y \sim p(y|\theta)$ an acceptance criterion $K_{\varepsilon_t}(D|y)$ is checked, with $t_1 > \dots, t_{n_t} \geq 0$. Generation $t$ returns a population of weighted parameters $\{(\theta_i^t, w_i^t)\}_{i\leq N} \sim \pi_{\text{ABC},\varepsilon_t}\theta|D)$ with $w(\theta) := \frac{\pi(\theta)}{g_t(\theta)}$. Usually, we are only interested in the last generation's particles.

*Remark* 6. While we here still have normalized importance weights, as soon as we work with non-normalized density functions later, we need to normalize the weights, using the biased estimator $W_i := w_i / \sum_{i=1}^N w_i$ before approximating integrals.

##### 3.2 An exact sequential ABC sampler

To make it more efficient and robust, we want to integrate the sampler introduced in Section 2 in a sequential Monte Carlo scheme. Based on ideas from parallel tempering and classical likelihood-based SMC, in generation $t$ we sample from

$$\pi_{\text{ABC},t}(\theta|D) \propto \int \pi(D|y,\theta)^{1/T_t} p(y|\theta)\, dy \cdot \pi(\theta), \tag{4}$$

where $T_1 > \ldots > T_{n_t} = 1$ are temperatures bridging from sampling from the prior to sampling from the posterior. Theorem 2 tells that for $T = 1$, we have exact inference from the correct posterior, i.e. with the correct assumption that the data are a realization of the model and additional measurement noise.

With some normalization constant $c_t$, the acceptance step in generation $t$ is given as

3. accept with probability $\left(\frac{\pi(D|y,\theta)}{c_t}\right)^{1/T_t}$.

##### 3.3 Choose the normalization constant automatically

In general, it is difficult to find a tight bound $c$ yielding high acceptance rates. For deterministic models, a value can be determined by a prior optimization, but in general this is not applicable. We propose to automatically select and adjust $c$ in a sequential scheme as follows: After iteration $t$, we set the value $c_{t+1}$ to the maximum value $\pi(D|y_i, \theta_i)$ sampled so far. Before the first iteration, we generate a calibration sample from the prior to obtain an initial guess $c_1$.

Now it can and in practice frequently does happen, because sampling tends to concentrate in high density regions, that in generation $t$ a value $\pi(D|y, \theta) > c_t$ is encountered. This means that the assumptions in Theorem 2 are violated and we no longer sample from the correct posterior. Figuratively, the acceptance probability of such particles is too small relative to particles in lower density regions, creating a bias. It is straightforward to imagine that we can correct for this bias by assigning a higher weight to particles violating the normalization constraint. That this indeed works out is content of the following theorem.

**Theorem 3** (Importance weighted acceptance)**.** *Consider a prior density $\pi(\theta)$, a proposal density $g_t(\theta)$, a model likelihood $p(y|\theta)$, a noise model $\pi(\bar{y}|y,\theta)$, let $D \sim q(\bar{y}|\theta) \propto\sim \pi(\bar{y}|y,\theta)p(y|\theta)\, dy$, and $T_t > 0$, $c_t > 0$ arbitrary. Then if we define the acceptance step to be*

3. accept with probability $\min\left[\left(\frac{\pi(D|y,\theta)}{c_t}\right)^{1/T_t}, 1\right]$

*and modify the importance weights to be*

$$w(y,\theta) \propto \frac{\pi(D|y,\theta)^{1/T_t}}{\min\left[\left(\frac{\pi(D|y,\theta)}{c_t}\right)^{1/T_t}, 1\right]} \cdot \frac{\pi(\theta)}{g_t(\theta)}, \tag{5}$$

*the weighted samples $(\theta, w(\theta))$ are sampled from $\pi_{\text{ABC},t}(\theta|D) \propto \int \pi(D|y,\theta)^{1/T_t} \pi(y|\theta)\, dy \cdot \pi(\theta)$. In particular, for $T = 1$ we sample from the correct posterior.*

*Proof.* Similar to the proof of Theorem 2, with the acceptance event $A = I[(y,\theta) \text{ is accepted}]$, we draw samples in joint space from

$$\pi(y,\theta|A=1) \propto \min\left[\left(\frac{\pi(D|y,\theta)}{c_t}\right)^{1/T_t}, 1\right] p(y|\theta) g_t(\theta),$$

while our target distribution remains

$$\pi_{\mathrm{ABC},t}(y,\theta|D) \propto \pi(D|y,\theta)^{1/T_t} p(y|\theta)\pi(\theta).$$

Thus, we need to apply exactly the importance weights (5), and obtain samples for $\theta$ alone by marginalization. □

In this study, we by default set $c$ to the maximum of the values found in previous iterations, which keeps differences in the weights as low as possible. When acceptance rates turn out too low, we set it to $\beta^{-T}c$, where $c$ is the maximum found value, and $\beta \geq 1$ increases acceptance rates roughly by that factor in the next iteration. Other schemes, e.g. based on quantiles, are possible. Before iteration 1, we draw a calibration sample from the prior.

#### 3.4 How to choose temperatures

We propose one class of geometric progression based schemes, and a scheme targeting acceptance rates, as detailed in the following.

##### 3.4.1 Choose the temperature to match a target acceptance rate

The idea of this scheme is to match a specified target acceptance rate, i.e. to choose $T_t = T$ s.t. the expected acceptance rate

$$\begin{aligned}\gamma &= \int \left( \int \min\left[ \left( \frac{\pi(D|y,\theta)}{c_t} \right)^{1/T}, 1 \right] p(y|\theta)\, dy \right) g_t(\theta)\, d\theta \\ &= \int w_t(\theta) \left( \int \min\left[ \left( \frac{\pi(D|y,\theta)}{c_t} \right)^{1/T}, 1 \right] p(y|\theta)\, dy \right) g_{t-1}(\theta)\, d\theta \\ &\approx \frac{1}{N} \sum_{i=1}^{N} w_t(\theta_i^{(t-1)}) \min\left[ \left( \frac{\pi(D|y_i^{(t-1)}, \theta_i^{(t-1)})}{c_t} \right)^{1/T}, 1 \right]\end{aligned} \tag{6}$$

matches a specified target rate. Here, in the second line we employ importance sampling from the previous proposal distribution $g_{t-1}$ with corresponding Radon-Nikodym derivatives $w_t(\theta) = g_t(\theta)/g_{t-1}(\theta)$. This is because in the third line, we approximate this integral via a Monte Carlo sample, using all parameters sampled in the previous iteration. Note: This must include also rejected particles to avoid a bias. The inner integral is approximated by the corresponding single simulation. The Radon-Nikodym derivatives can be omitted if subsequent proposal distributions are sufficiently close. In practice, it may suffice to consider a subset of all particles to accelerate computations.

Matching $\gamma \approx \gamma_{\text{target}}$ (we usually used $\gamma_{\text{target}} = 0.3$) is a simple one-dimensional bounded optimization problem. It should be noted that this scheme, here referred to as *acceptance rate scheme*, is just a rough heuristic estimate of the expected acceptance rate. However, for our purpose proved sufficient.

In particular, this scheme can also be used for the first iteration, if a calibration sample from the prior is employed, in which case $g_{t-1}(\theta) = g_t(\theta) = \pi(\theta)$. But also in later iterations it may be useful, if other schemes propose too hesitant a decrease, while an acceptance rate of $\gamma$ is usually justifiable. In particular, we observed huge leaps to be possible in the second iteration.

In later iterations however, it may be impossible to maintain high acceptance rates without decreasing the $c$ value. Inserting for $g(\theta)$ either the prior $\pi(\theta)$ or the posterior $\pi(\theta|D)$ allows for the comparison of those two theoretical acceptance rates. Sampling from the posterior will usually lead to a higher acceptance rate, since it is more concentrated around the observed data, in particular if the data are informative of the parameters. $\gamma_{\pi(\theta|D)}$ is expected to be close to the acceptance rate achieved in the later iterations of an ABC-SMC sampler, when the proposal distribution is already quite close to the posterior, while simple ABC-Rejection from the prior gives a theoretical acceptance rate of $\gamma_{\pi(\theta)}$. However, $\gamma_{\pi(\theta|D)}$ can be arbitrarily small, as it depends on the model and the data. Thus, other methods for reducing the temperature need to be applied then, forcing it down.

###### 3.4.2 Update temperature in an exponential decay

In standard likelihood-based Markov Chain Monte Carlo (MCMC) sampling, empirically a geometric progression has shown to yield equal probabilities for swaps between adjacent temperatures Sugita et al. [2000]. That is a scheme with fixed temperature ratios, thus linear in log-space. This finding can be theoretically justified under some conditions Predescu et al. [2004]. Since a similar approach was recently successfully applied in an ABC-SMC setting Daly et al. [2017], we chose to use a geometric progression here too, referred to as *exponential decay scheme*. Either we specify a fixed ratio $\alpha \in (0, 1)$ s.t.

$$T_{t+1} = \alpha T_t$$

(we used by default a value $\alpha = 0.5$), or we pre-select the number of iterations $N$ and accordingly set

$$T_{t+1} = T_t^{(N-t)/(N-t+1)} = T_1^{(N-t)/N},$$

where typical numbers of iterations $N$ in ABC-SMC are 10 to 30. In this study, we only employed the former scheme. It should be noted that the underlying motivation is very heuristic, since the SMC setup is different from MCMC. There exist approaches to dynamically adjust the temperatures Predescu et al. [2004], Vousden et al. [2015], however these are specific to MCMC settings. For likelihood based SMC, there exist approaches that try to keep the effective sample size constant Latz et al. [2018], however it remains to be investigated whether these are applicable in a likelihood-free context, which goes beyond the present work.

Assuming both schemes to deliver appropriate proposals, however possibly too hesitant, we set the effective temperature used in the next iteration as the minimum of the values proposed by the different schemes.

For brevity, we denote by ASSA in the following the here proposed exact ABC-SMC sampler with an adaptive sequential stochastic acceptor, i.e. with $c$ set to the previously observed highest value with weight correction, and the two temperature selection schemes, with target acceptance rate $\gamma_{\text{target}} = 0.3$, and $\alpha = 0.5$ in the exponential decay scheme.

### 4 Upper bounds on the normalization constant

When not employing a self-tuned routine to find good values for the normalization constant $c$, and without reweighting, a tight upper bound on $(y, \theta) \mapsto c = \pi(D|y, \theta)$ must be identified in order to keep acceptance rates reasonable. Without further information on the model, the only feasible way is to set it to $\hat{c} = \max_{y,\theta} \pi(D|y, \theta)$, where $y$ can take arbitrary values, not confined to realization possible under the model.

#### 4.1 Normal and Laplace noise

Assuming the noise parameters, i.e. respectively for normal and Laplace noise the standard deviations $\sigma$ and the scale parameters $b$, to be fixed, for the normal and Laplace distribution the highest peak of $y \mapsto \pi(D|y)$ ($= \pi(D|y, \theta) \forall \theta$) is at the mode of the distributions, i.e. at $y = D$. If in a normal or Laplace noise model the noise parameters are also estimated as part of $\theta$, then the maximum value of $\pi(D|y, \theta)$ is achieved for minimum $\sigma, b$.

#### 4.2 Poisson noise

For the Poisson distribution, where the variance depends on the simulation, the same holds, as we shortly verify: The distribution is given as

$$f(n,k) = \frac{n^k}{k!}e^{-n},$$

assuming $n$ to be the simulated value and $k$ the noisy data. In our application, thus $n$ is the latent real noise-free value of mRNAs, and $k$ is the experimentally measured value. To find the mode of this function over $n$ (not over the distribution parameter $k$, because we need an upper bound over all model simulations $n$ while keeping $k$ fixed), we differentiate

$$\nabla_n f(n,k) = (kn^{k-1} - n^k)e^{-n} > 0 \Leftrightarrow kn^{k-1} > n^k \Leftrightarrow n < k,$$

i.e. the mode is indeed at $n = k$.

#### 4.3 Binomial noise

In addition, let us shortly derive the solution assuming a binomial noise model (implemented, but not used in the study): The function is

$$f(n,k) = \binom{n}{k} p^k (1-p)^{n-k} \tag{7}$$

for a success probability $p \in [0,1]$, number of experiments $n$ and number of successes $k$. We compute the ratio

$$\frac{f(n+1,k)}{f(n,k)} = \frac{\binom{n+1}{k} p^k(1-p)^{n+1-k}}{\binom{n}{k} p^k(1-p)^{n-k}} = \frac{(n+1)(1-p)}{n+1-k} < 1 \Leftrightarrow -np - p < -k \Leftrightarrow n > \frac{1}{p}(k-p).$$

Thus, the modal value is at $n = \lceil \frac{1}{p}(k-p) \rceil$ or $n = \lceil \frac{1}{p}(k-p) \rceil + 1$.

### 5 Implementation

#### 5.1 Implementation in the toolbox pyABC

All of the discussed methods have been implemented in the open-source toolbox pyABC (`https://github.com/icb-dcm/pyabc`) and are thus directly accessible to the community. pyABC implements an ABC-SMC algorithm based on Toni et al. [2008] with various extensions. In particular, it comes directly with high performance computing capabilities, which we also exploited in our analysis. It is written in python but supports models in arbitrary languages. We put special emphasis on an easy-to-use modular implementation, s.t. it is straightforward to customize the analysis pipeline.

pyABC has been developed with a classical ABC setup in mind, i.e. using a distance and an epsilon threshold. To use the here presented noisy approach, one can proceed as follows: Instead of a distance, define the noise likelihood e.g. via a `NormalKernel` for a normal noise model, but also other kernels are implemented. Instead of an epsilon threshold, define a temperature. Then put both together in an acceptance step. In code, this looks like:

```
kernel = pyabc.NormalKernel()
temperature = pyabc.Temperature()
acceptor = pyabc.StochasticAcceptor()
```

All of these classes take various configuration arguments, but already the defaults have been shown to be quite stable. In particular, the temperature scheme is set to use both the acceptance rate schedule and a geometric progression, and the acceptor uses the adaptive normalization approach.

Given a noise-free `model` and a `prior`, some storage location `database` and observed data `observation`, then the analysis can be started as simple as:

```
abc = pyabc.ABCSMC(models=model,
                   parameter_priors=prior,
                   distance_function=kernel,
                   eps=temperature,
                   acceptor=acceptor)
abc.new(db=database,
        observed_sum_stat=observation)
abc.run()
```

#### 5.2 Numerical considerations

From an implementation point of view, it is temperature ratios that are observed to be effective, such that we solve the acceptance rate prediction problem for $\log T \in (-\infty, 0]$ on a log-scale, where for numerical purposes we use a lower boundary $-100 \leq \log T$. This boundary was however never hit in practice. Since the computational effort scales linearly in the number of simulations, we allow to limit the number of simulations recorded via the `pyabc.ABCSMC.max_nr_recorded_particles` parameter, e.g. to 10 times the population size. Since we are interested in the application to expensive multi-scale models, whose simulation effort overwhelms, we did not consider this here.

As the likelihood decreases exponentially with the number of data points, we also perform the temperature exponentiation in log space.

#### 5.3 Computational resources

We ran the analyses on three types of hardware: Smaller problems were run on a non-dedicated desktop computer with 4 Intel Core i7-6500U CPUs (2.50GHz) and 16GB RAM. Medium-scale problems M1-5 were run on a non-dedicated computer with 48 Intel Xeon Gold 6125 cores (2.60GHz) and 277GB RAM, using up to 40 cores. The computationally expensive problem M6 was run on a computational cluster, using up to 200 cores. As we used the number of samples as a measure of efficiency, the exact specifications of the hardware are of no consequence. For the analyses on a single computer, the pyABC `pyabc.sampler.MulticoreEvalParallelSampler` was used, using multiprocessing, for the analyses on the cluster the `pyabc.sampler.RedisEvalParallelSampler` was used, enabling distributed processing.

#### 5.4 Code and data

The entire code and data underlying this study have been uploaded to zenodo at http://doi.org/10.5281/zenodo.3631120. The code alone, without the databases, is also available on GitHub: https://github.com/yannikschaelte/Study-ABC-Noise.

### 6 Examples of ignoring measurement noise

#### 6.1 The ODE model from the main manuscript

The ordinary differential equation (ODE) model presented in the main manuscript is model M1 of a conversion reaction, confined to parameter $\theta_1$, with 10 data points.

#### 6.2 A non-identifiable model

While for the model disucssed in the main manuscript we obtain point estimates, this does not need to be the case, even for deterministic models. For example, consider an ordinary differential equation (ODE) model of the simplistic reaction system

$$A \xrightarrow{p_0} \emptyset, \quad A \xrightarrow{p_1} \emptyset,$$

i.e. a speces decaying at two different rates. The analytical solution is

$$A(t) = \exp((p_0 - p_1)t)A_0,$$

i.e. only the difference $p_0 - p_1$ is identifiable. For true parameters $\theta_{\text{true}} = (p_0, p_1) = (0.4, 0.5)$ and $A_0 = 1$, we assumed as observables 6 equally spaced measurements in the interval $[0, 30]$. We simulated data from the true parameters and created artificial noisy data by adding independent normal noise of standard variation $\sigma = 0.2$ (Figure S1A).

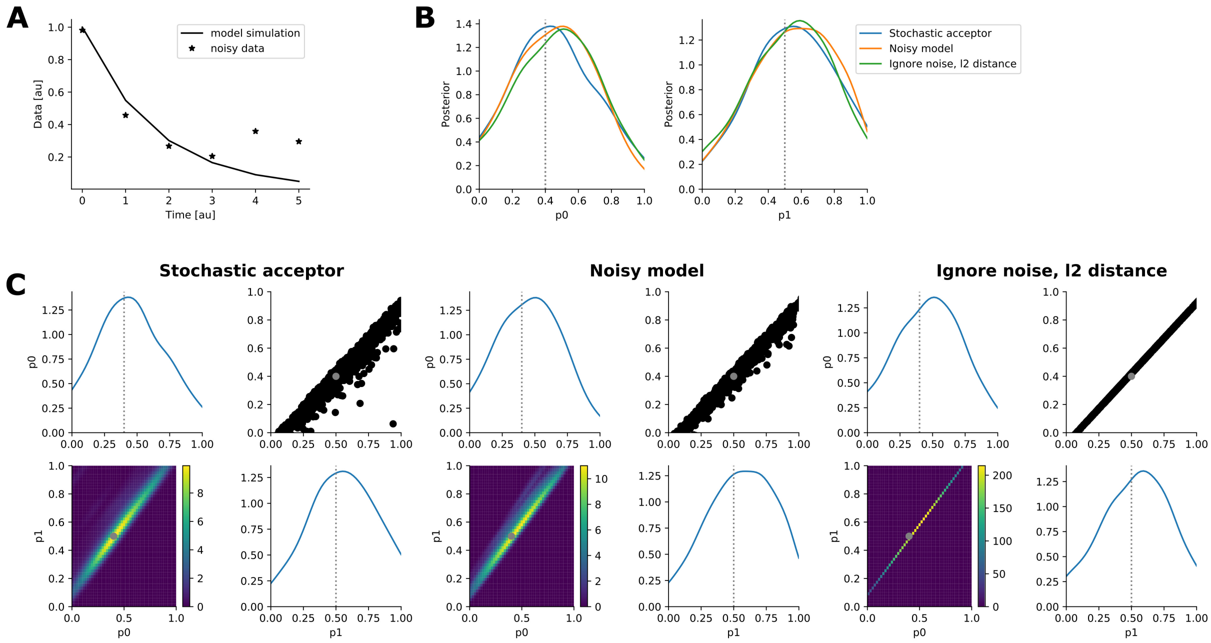

Figure S1: Illustrations for the non-identifiable motivational model. (A) The noise-free model simulation and the noisy data. (B) Kernel density estimates (KDE) of the 1-dimensional posterior marginals for all three employed ABC analyses. (C) KDEs of the 1- and 2-dimensional posterior margins for the three employed analyses. In each 2-by-2 plot, on the top left and bottom right the 1-dimensional marginals are shown, on the bottom left the 2-dimensional distribution visualized via a heat map, and on the top right the actual $N = 1000$ samples illustrated as black dots. In (B) and (C) the dotted grey lines indicate the true parameter values.

Assuming a flat prior over $[0, 1]$ for both parameters, we performed ABC inference (I) employing the stochastic acceptor introduced in the main manuscript as A3 in the optimized version developed in the main manuscript,

(II) randomizing the model output by adding the same normal distribution, and (III) using the noise-free model and completely ignoring the noise by applying standard ABC with the threshold $\varepsilon$ decreased as far as the computational budget allowed, and using an $\ell_2$ distance corresponding.

The 1-dimensional posterior marginals generated by the three approaches are not distinguishable (Figure S1B). In particular, compared to the model discussed in the main manuscript, they do not converge to point estimates, because there cannot be a single optimal parameter, but only an optimal difference $p_0 - p_1$. However, the 2-dimensional marginals differ (Figure S1C). Here, it becomes apparent that approach (III) yields a degenerate distribution, essentially only a line, which in addition does not capture the true parameters. In contrast, the stochastic acceptor (I) yields a reasonable non-degenerate posterior distribution. The noisy model (II) yields the same distribution, confirming the result.

This illustrates that even for deterministic models, the resulting posterior does not have to converge to a point estimate. Non-trivial distributions are possible, if the parameters are not completely identifiable.

#### 6.3 A stochastic differential equation model

If we apply standard ABC to deterministic mechanistic models with noisy data, we are usually not able to reproduce the data exactly and thus encounter a minimum threshold $\varepsilon_{\min}$ and convergence to points $y = y(\theta)$ possessing that exact distance value. If the mechanistic model $p(y|\theta)$ is itself inherently stochastic, even when not accounting for model error in general a non-degenerate distribution is to be expected, because for the same parameters, the model does not always return the same simulations.

To illustrate how measurement noise can lead to wrong parameter estimates for stochastic models, we firstly employ the stochastic differential equation (SDE) model M3 from the main manuscript (described in detail in the following chapter), with prior boundaries of dc $\in [18, 25]$, membrane_dim $\in [5, 15]$.

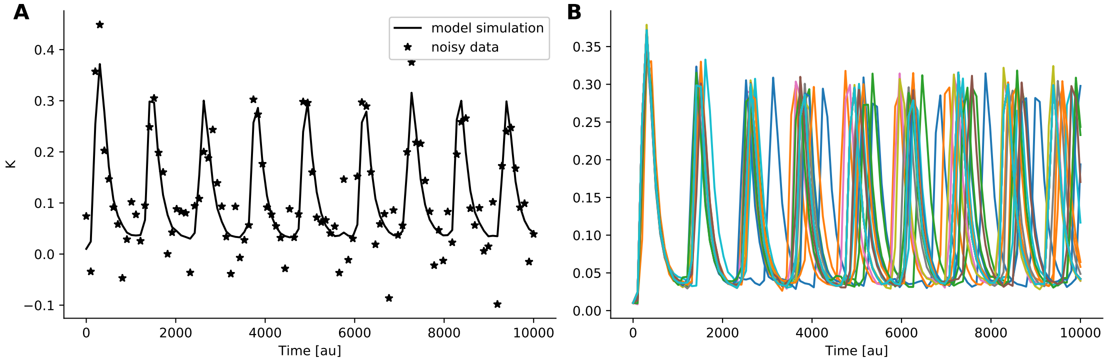

Figure S2: Model simulations and data for the SDE motivational model. (A) The model simulation and noisy data employed in this analysis. (B) 20 runs of the model, as an illustration of the degree of inherent stochasticity.

We assumed an additive normal noise model and generated the data accordingly (Figure S2). Then, we performed three analyses: First, we ignored the noise and performed standard ABC with an $\ell_2$ distance. Second, we performed correct exact inference using the stochastic acceptor. Third, we also performed inference using standard ABD and an $\ell_2$ distance, but for the non-noisy data.

The run operating on the noise-free data was able to correctly and succinctly identify the true parameters (Figure S3A). Of the two runs operating on the noisy data, the one ignoring noise yielded a very succinct posterior distribution, close to a point estimate, however not at the true parameters (Figure S3A). The run accounting for noise shows that $\text{membrane}_\text{d}\text{im}$ could not be well identified, for dc the posterior was still confined. This situation is similar to the one obtained for the model shown in the main manuscript, despite the model stochasticity. In fact, for this model the run operating on the noisy data but ignoring noise the $\varepsilon$

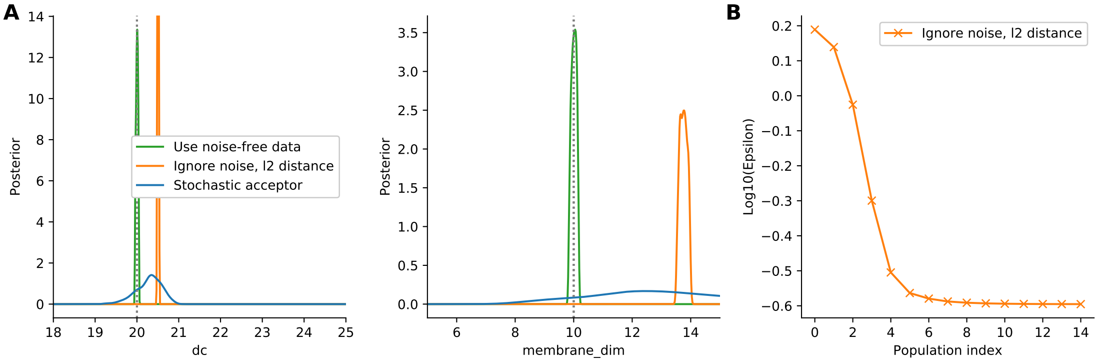

Figure S3: Analysis of the SDE motivational model. (A) KDEs of the posterior approximations obtained using the three setups. The runs “Ignore noise” and “Stochastic acceptor” are on the noisy data, “Use noise-free data” operates on the non-noisy model output. (B) The $\varepsilon$ thresholds employed by the run ignoring noise but using noisy data.

seemed to converge to a value greater than zero, corresponding to the stochastic model not being completely able to reproduce the data (Figure S3B).

#### 6.4 A Markov jump process model

As a second stochastic model, we investigated the impact of ignoring measurement noise on the Markov jump process (MJP) model M4 from the main manuscript, described in detail in the following chapter, confined to one parameter transctiption. For this model, we assumed a Poisson noise model, a common assumption for count data (Figure S4A). The simulations produced by this model possess a high variability (Figure S4B) and thus are not unlikely to be able to produce the noisy data. Thus the results obtained from an analysis may be expected to differ from those obtained for model M3.

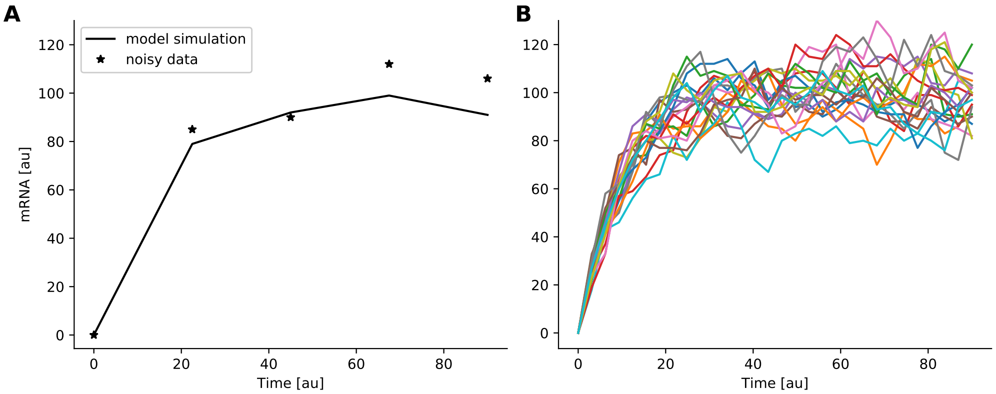

Figure S4: Model simulations and data for the MJP motivational model. (A) The model simulation and noisy data employed in this analysis. (B) 20 runs of the model, as an illustration of the degree of inherent stochasticity. The trajectories shown here have a higher resolution than the 5 data points, to capture the fluctuations of this model on short time scales.

Again, we performed three analyses: First, we ignored the noise and performed standard ABC with an $\ell_2$ distance. Second, we performed correct exact inference using the stochastic acceptor. Third, we also performed inference using standard ABC and an $\ell_2$ distance, but for the non-noisy data.

Indeed, for this model the posterior approximations obtained using all three approaches are non-degenerate, even assuming noise-free data (Figure S5A). This indicates that the randomness of the model itself is com-

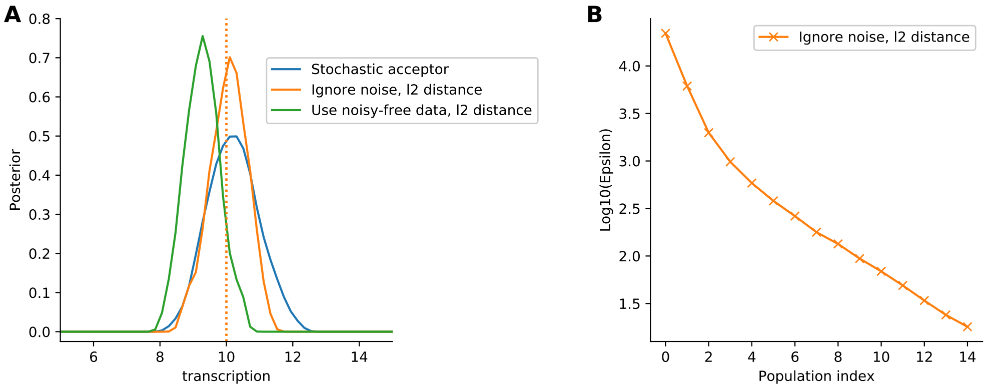

Figure S5: Analysis of the MJP motivational model. (A) KDEs of the posterior approximations obtained using the three setups. The runs "Ignore noise" and "Stochastic acceptor" are on the noisy data, "Use noise-free data" operates on the non-noisy model output. (B) The $\varepsilon$ thresholds employed by the run ignoring noise but using noisy data.

parably high, allowing various parameter configurations to produce data matching the observed data closely. This fits to no convergence to a value greater than 0 being observed for $\varepsilon$ (Figure S5 B) at the point where the analysis was stopped due to a too low acceptance rate. The difference between approximations for the noisy data is not as big as for previous models. If measurement noise is correctly accounted for, the distribution becomes wider (Figure S5A), but both are similarly centered around the true parameters. One explanation of this is that for this example, the heterogeneity of the model is relatively large compared to the impact of the noise model.

#### 6.5 A Markov jump process model with a non-centered noise model

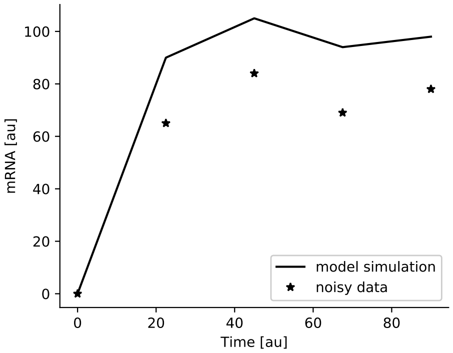

Figure S6: Data for the MJP motivational model with binomial noise.

The situation considered in the previous section changes if we, just for the sake of the example, consider an alternative noise model. In this section, we assume, instead of the Poisson noise, a binomial noise model. A motivation of this noise could be that e.g. mRNAs or proteins can only be detected with a certain accuracy in fluorescence microscopy.

Supposing a detection rate of only 0.7, the simulated data points are visibly below the noise-free simulations (Figure S6. Applying standard ABC ignoring noise, in this situation the inferred transcription rate parameter is below the true one (Figure S7A). But if we correct for the measurement noise by performing exact inference under the assumption of a binomial noise model using the stochastic acceptor, the posterior curve is shifted back to higher values.

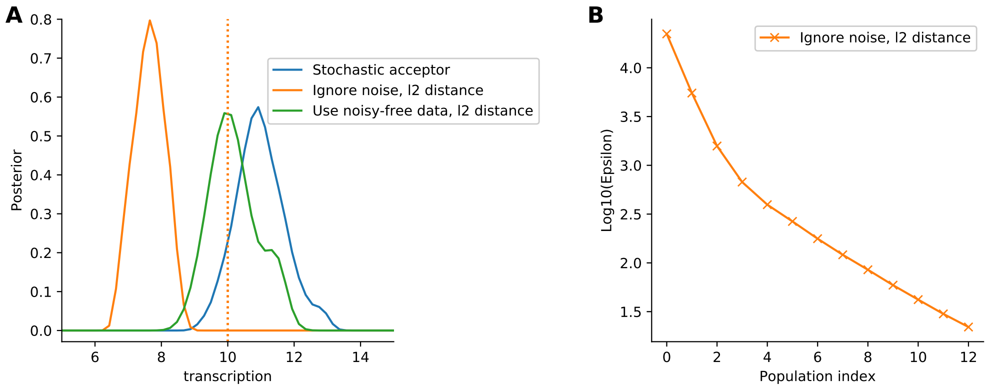

Figure S7: Analysis of the MJP motivational model with binomial noise. (A) KDEs of the posterior approximations obtained using the three setups. The runs “Ignore noise” and “Stochastic acceptor” are on the noisy data, “Use noise-free data” operates on the non-noisy model output. (B) The $\varepsilon$ thresholds employed by the run ignoring noise but using noisy data.

### 7 Details on the test models

In this chapter, the test models employed in the results part of the main manuscript are explained in further detail.

#### 7.1 M1 and M2: Conversion reaction

Model M1 is an ODE model of a simple conversion reaction system,

$$A \underset{\theta_2}{\overset{\theta_1}{\rightleftharpoons}} B,$$

the analytical solution to which is

$$\begin{pmatrix} A \\ B \end{pmatrix} (t, \theta) = \frac{1}{\theta_1 + \theta_2} \left[ \begin{pmatrix} \theta_2 & \theta_2 \\ \theta_1 & \theta_1 \end{pmatrix} - \begin{pmatrix} -\theta_1 & \theta_2 \\ \theta_1 & -\theta_2 \end{pmatrix} \exp(-(\theta_1 + \theta_2)t) \right] \begin{pmatrix} A_0 \\ B_0 \end{pmatrix}$$

with parameter vector $\theta = (\theta_1, \theta_2)$. We assumed initial concentrations $(A_0, B_0) = (1, 0)$ and only species $A$ to be measured. Both reaction rate constants were estimated. In the standard estimation problem, we assumed to have $n_y = 10$ equidistant measurement points in the interval $[0, 30]$. The measured data were assumed to be noise-corrupted by additive normal noise with a standard deviation of $\sigma = 0.02$. We thus created synthetic data as $D \sim (A(t_1, \theta), \dots, A(t_m, \theta)) + \mathcal{N}(0, \sigma I_{n_y})$, assuming the true parameters to be $\theta_{\text{true}} = (\theta_1, \theta_2) = (0.06, 0.08)$. We imposed an uninformative uniform prior over $[0, 0.4]$ for both parameters. As distance for the noisy model approach we employed a weighted $\ell_2$ distance $d(x, y) = \sum_{i=1}^{m} \omega_i (x_i - y_i)^2$ with $\omega_i = \sigma^{-1}$ for all $i$, while for the stochastic acceptor approaches the acceptance step was based on the true normal distribution.

As it is frequently employed to model outlier-corrupted data, we also considered model M1 with an additive Laplace noise model with scale parameter $b = 0.02$. As distance measure for the noisy model approach, the $\ell_2$ distance was further used, while the kernel for the stochastic acceptor was adapted.

#### 7.2 M3: Hodgkin-Huxley neurons

Model M3 is a stochastic differential equation (SDE) model of intrinsic ion channel noise in Hodgkin-Huxley neurons, based on Goldwyn et al. [2011]. We used a Fortran 95 implementation made available on ModelDB (https://senselab.med.yale.edu/ModelDB) under Accession 128502. We assumed only the fraction of open $K$ channels to be measured at $n_y = 100$ equidistant time steps over a time frame covering roughly 9 oscillation periods of the model, and employed an additive normal noise model of standard deviation $\sigma = 0.05$ to describe inaccuracies in the measurement process. We estimated the parameters dc describing the input current, and the square root of the membrane area membrane_dim. We assumed uniform priors $\text{dc} \in [2, 30], \text{membrane\_dim} \in [1, 12]$, with ground truth parameters $(\text{dc}_{\text{true}}, \text{membrane\_dim}_{\text{true}}) = (20, 10)$.

#### 7.3 M4: mRNA transcription

Model M4 is a stochastic model describing the process of mRNA transcription, $\emptyset \overset{p_1}{\to} \text{mRNA} \overset{p_2}{\to} \emptyset$, with $\text{mRNA}_0 = 0$. We estimated the transcription rate constant $p_1$ and the decay rate constant $p_2$, assuming a uniform prior over $p_1 \in [0, 30], p_2 \in [0, 0.2]$, and ground truth parameter $p_{1,\text{true}} = 10, p_{2,\text{true}} = 0.1$. To capture the intrinsic stochasticity of this process at low copy numbers, we sampled from the chemical master equation using the Gillespie algorithm. We assumed mRNA counts from a single cell to be available at $n_y = 10$ time points in the interval $[0, 90]$. We employed a Poisson noise model, which is frequently used for regression of count data.

##### 7.4 M5: STAT5 dimerization

Model M5 is an ODE model by Boehm et al. [2014] describing the homo- and heterodimerization of the transcription factors STAT5A and STAT5B. The model is defined in SBML Hucka et al. [2003] and taken from the systems biology benchmark collection Hass et al. [2019]. The model and data were converted to the PEtab format Weindl et al. [2019] (version 0.0.0a13), and the tool AMICI Fröhlich et al. [2017] (version 0.10.13) used to efficiently simulate the ODE solutions numerically. As in the original publication, uniform parameter priors and additive normal measurement noise were assumed. We estimated 11 logarithmically scaled parameters of this model, including three standard deviations of the normal noise model, one for each data type.

##### 7.5 M6: Tumor spheroid growth

Model M6 is a multi-scale ABM model of spheroid tumor growth on a two-dimensional plane, as described in Jagiella et al. [2017]. The model implementation is in C++ with Python bindings and can be found at https://github.com/ICB-DCM/tumor2d. Up to roughly a million single cells were modeled as stochastically interacting agents, coupled to the dynamics of extracellular substances modeled via partial differential equations. Data produced by this model are the spheroid radius over time, and, at discrete intervals from the spheroid rim, the fraction of proliferating cells and the extracellular matrix (ECM) intensity. For each of the three data types, 10 time-equidistant data points along the interesting part of the system dynamics were assumed to be available. An additive normal noise model was assumed, with standard deviations of $\sigma = 40, 0.15, 0.02$, respectively, for the three data types.

##### 7.6 Data

A selection of the data employed for models M1-6 in the study is shown in Figure S8.

#### 8 Additional analyses for the test models

##### 8.1 Comparison of inferred and theoretical posterior distribution for M1 and M2

The comparison of the posterior distributions computed by ASSA for models M1 and M2 with the theoretical posterior verifies that the algorithm yields the correction distribution (Figure S9).

##### 8.2 Temperature values proposed by the update schemes

In Figure S10, it is exemplarily shown how the acceptance rate scheme proposed an initial temperature, and then also in the first iteration proposed a value which was below the one suggested by the exponential decay scheme, thus improving efficiency by moving to lower temperatures more quickly if the acceptance rate allows it. Thereafter, the exponential decay scheme took over until the end. For more complex models, the acceptance rate scheme could often not propose temperatures below some $T_{\min} \gg 1$, so that only the exponential decay scheme ensured convergence to $T = 1$.

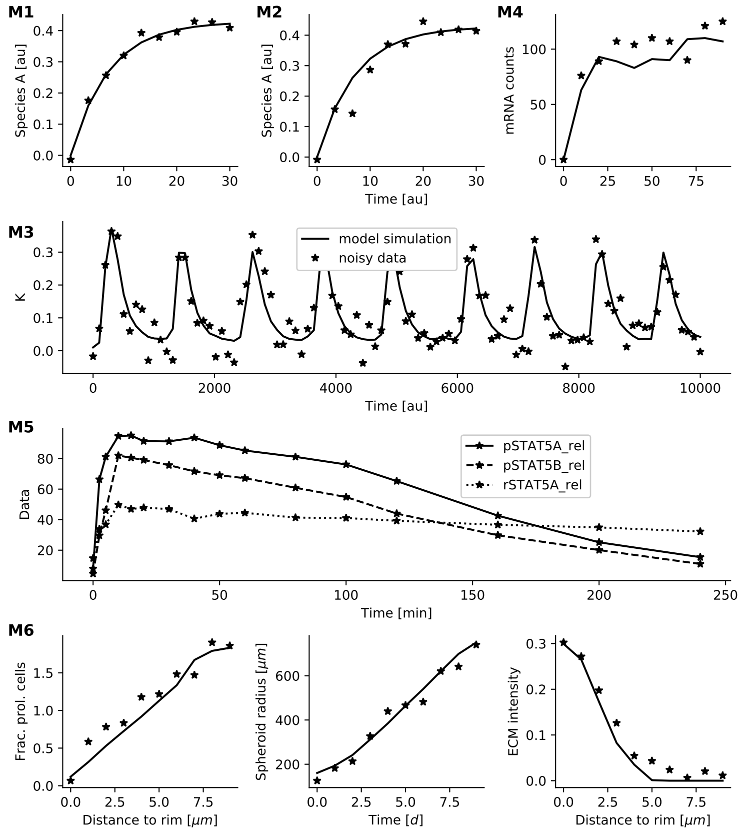

Figure S8: Selection of data sets employed in this study for the different models. For all but M5, ground truth simulations and noisy data created by adding noise to the simulations are shown. As for model M5, no ground truth exists, only the three data types are shown.

#### 8.3 Scaling study regarding the number of data points

In order to assess the influence of the number of data points on ASSA, for model M1-4 we altered the number of equidistant data points, leaving all other settings untouched. For M1-3, we chose numbers between 3 and 1000, for M4 between 3 and 15 due to the lower acceptance rates for this model.

As the number of data points increases exponentially for models M1-3, the number of simulations increases only moderately. This indicates that for these models, ASSA is fairly robust w.r.t. the number of data points. For model M4 however, the required number of simulations increased considerably. This could be explained by the high degree of stochasticity of this model, such that inference is in general hard.

Further, we looked at the ratios $\hat{c}/c$ for the different data points, where $\hat{c}$ denotes the, too high, upper limit on the noise likelihood when the ability of the model to generate such points is ignored, and $c$ is the upper limit experimentally found by ASSA. The ratio seems to grow exponentially with the number of data points for all of the models (Figure S12). This underlines the necessity to find good values for $c$.

#### 8.4 Effective sample size corrected by number of simulations

The effective sample sizes corrected by the number of simulations are shown in Figure S13.

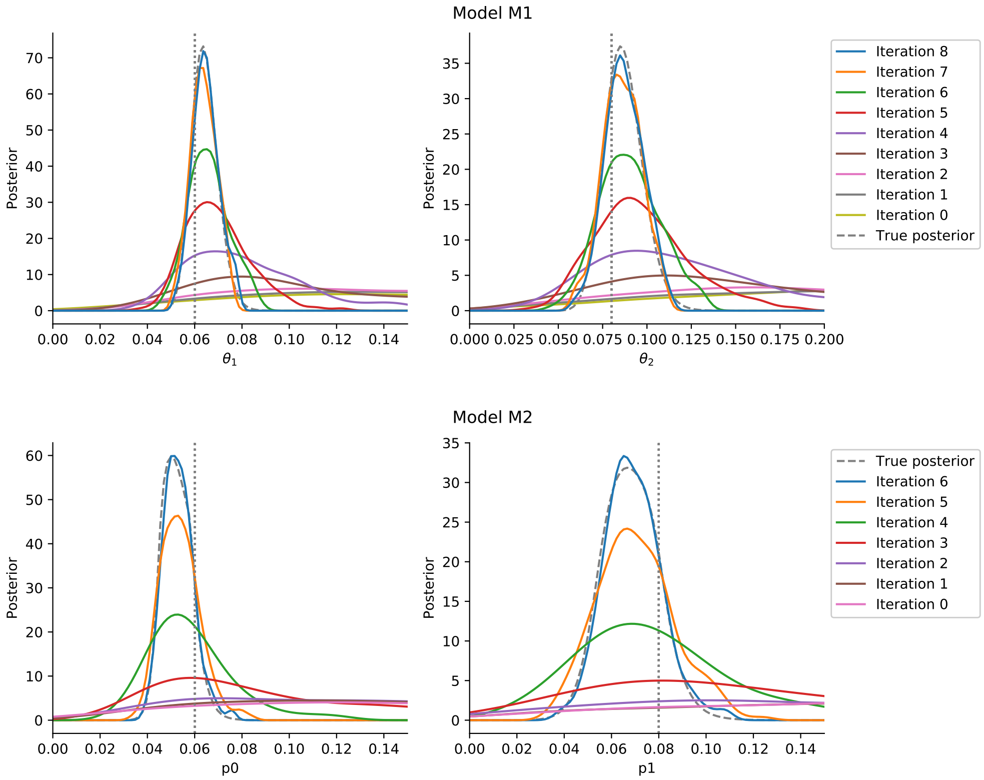

Figure S9: The posterior distributions inferred by ASSA for model M1 (top) and M2 (bottom) over the iterations, compared to the theoretical posterior distribution.

#### 8.5 M5: STAT5 dimerization

The data for the three observables corresponding to the parameters sampled in the last iteration of ASSA are plotted against the measured data in Figure S14. The fit is reasonable, the simulations closing following the observed data, but still possessing some variability, resulting from the assumed measurement noise model.

#### 8.6 M6: Tumor spheroid growth

The posterior marginals obtained using ASSA, and compared to values obtained using the noisy model sampler with a similar computational budget, are shown in Figure S15.

The data for the three observables corresponding to the parameters sampled in the last iteration of ASSA are plotted against the measured data in Figure S16. As for model M5, the fit is reasonable, closely matching but not degenerate.

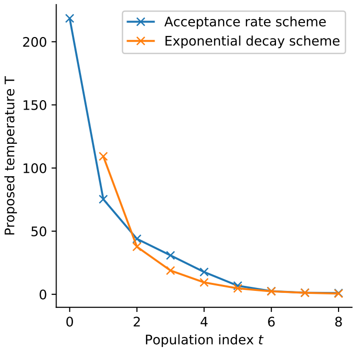

Figure S10: Temperatures chosen by the two update schemes for model M1. The value shown at iteration $t$ is based on the results of iteration $t-1$. The minimum proposed value was selected in each iteration.

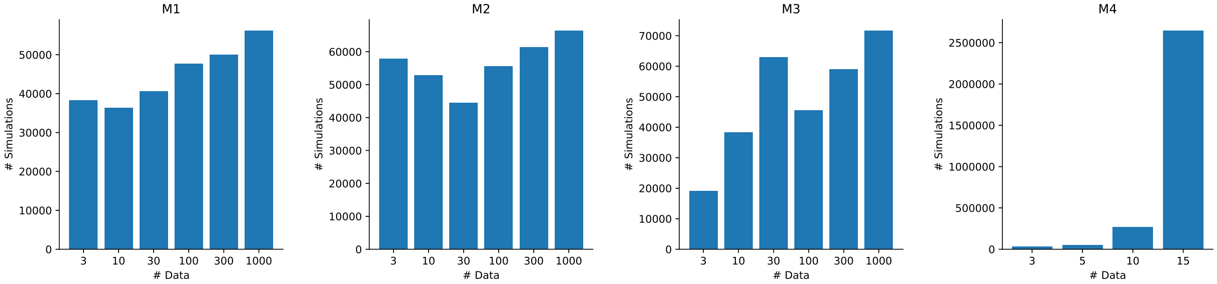

Figure S11: The numbers of simulations required by ASSA on the test models M1-4 for varying numbers of data points.

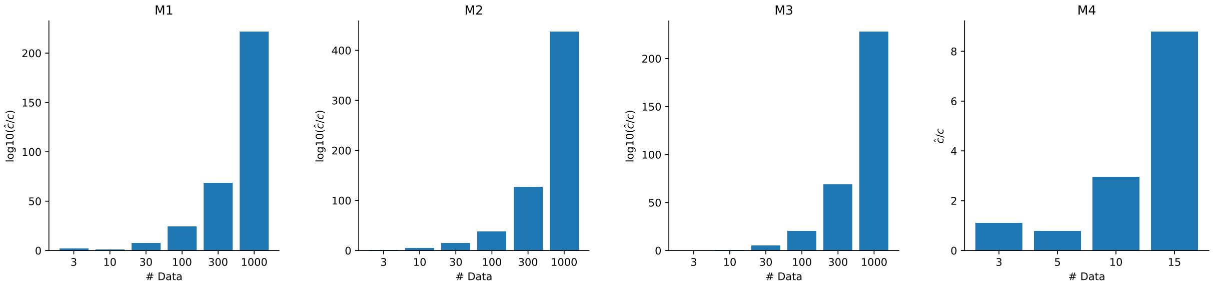

Figure S12: The factors $\hat{c}/c$ with $\hat{c}$ the theoretically obtained value for the highest peak of the noise model, and $c$ is the value found experimentally by ASSA. Note that the values for model M4 are plotted on a linear scale, the differences between data points are on a linear scale, not exponential as for the others.

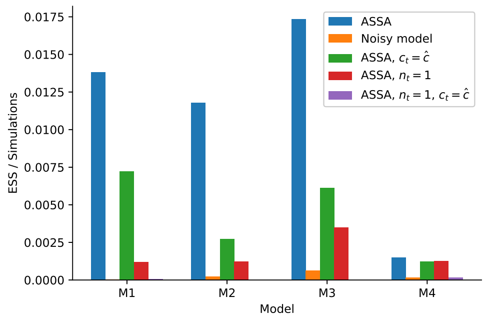

Figure S13: The effective sample sizes for the five samplers on models M1-4, divided by the total number of simulations. Note that for runs that did not stop with exact inference, $T = 1$, the values until reaching exact inference would be considerably smaller.

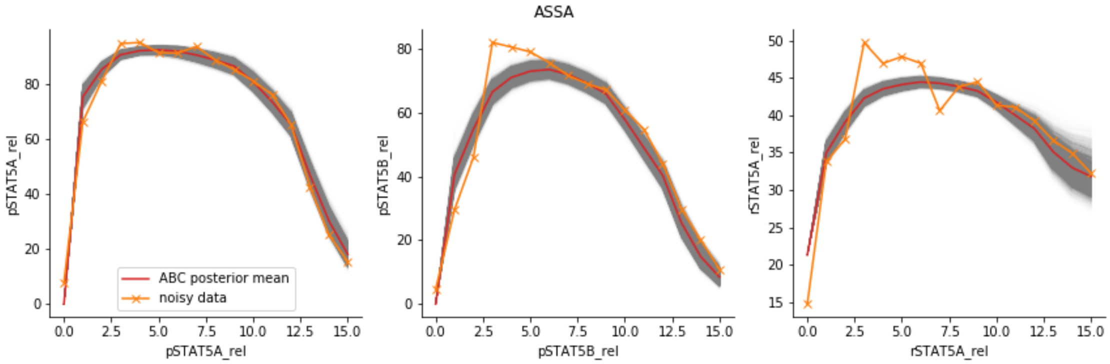

Figure S14: For model M5, the simulated data from the last iteration of ASSA for $N = 1e4$ simulations in grey, plotted against the measured data.

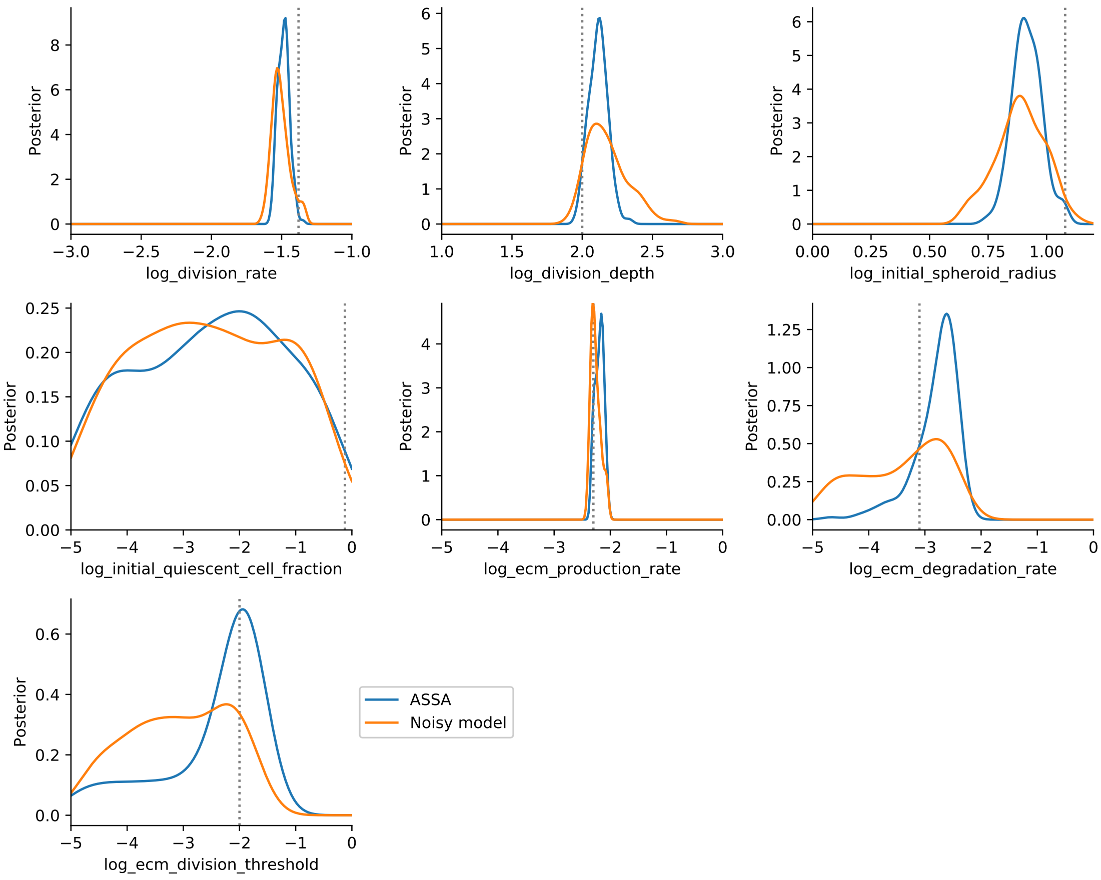

Figure S15: Posterior marginals for model M6, using ASSA and the noisy model sampler. Ground truth parameters are indicated by dotted lines.

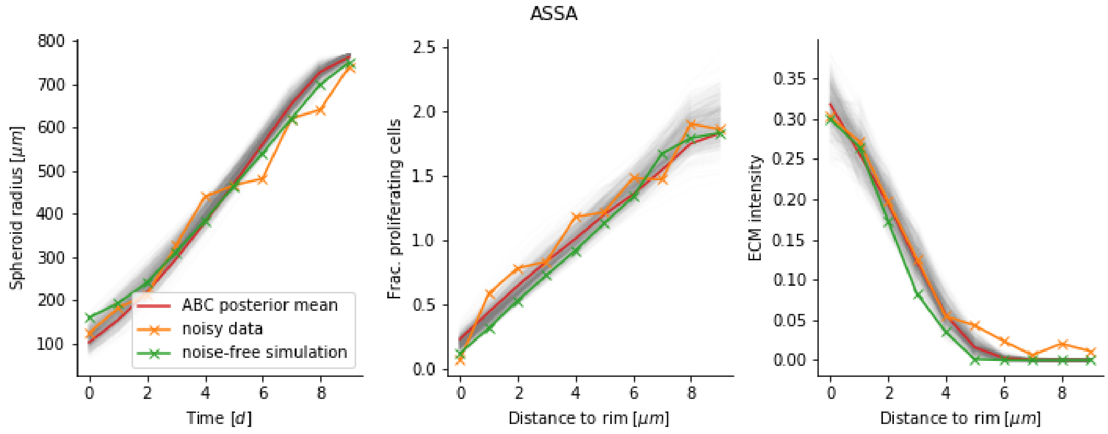

Figure S16: For model M6, the simulated data from the last iteration of ASSA for $N = 5e2$ simulations in grey, plotted against the measured data.

### References

Martin E Boehm, Lorenz Adlung, Marcel Schilling, Susanne Roth, Ursula Klingmueller, and Wolf D Lehmann. Identification of isoform-specific dynamics in phosphorylation-dependent stat5 dimerization by quantitative mass spectrometry and mathematical modeling. *Journal of proteome research*, 13(12):5685–5694, 2014.

Aidan C Daly, Jonathan Cooper, David J Gavaghan, and Chris Holmes. Comparing two sequential monte carlo samplers for exact and approximate bayesian inference on biological models. *Journal of The Royal Society Interface*, 14(134):20170340, 2017.

Fabian Fröhlich, Fabian J Theis, Joachim O Rädler, and Jan Hasenauer. Parameter estimation for dynamical systems with discrete events and logical operations. *Bioinformatics*, 33(7):1049–1056, 2017.

Joshua H Goldwyn, Nikita S Imennov, Michael Famulare, and Eric Shea-Brown. Stochastic differential equation models for ion channel noise in hodgkin-huxley neurons. *Physical Review E*, 83(4):041908, 2011.

Helge Hass, Carolin Loos, Elba Raimúndez-Álvarez, Jens Timmer, Jan Hasenauer, and Clemens Kreutz. Benchmark problems for dynamic modeling of intracellular processes. *Bioinformatics*, 35(17):3073–3082, 2019.

Michael Hucka, Andrew Finney, Herbert M Sauro, Hamid Bolouri, John C Doyle, Hiroaki Kitano, Adam P Arkin, Benjamin J Bornstein, Dennis Bray, Athel Cornish-Bowden, et al. The systems biology markup language (sbml): a medium for representation and exchange of biochemical network models. *Bioinformatics*, 19(4):524–531, 2003.

N. Jagiella, D. Rickert, F. J. Theis, and J. Hasenauer. Parallelization and high-performance computing enables automated statistical inference of multi-scale models. *Cell Systems*, 4(2):194–206, 2017.

Ram P Kanwal. *Generalized functions theory and technique: Theory and technique.* Springer Science & Business Media, 2012.

Jonas Latz, Iason Papaioannou, and Elisabeth Ullmann. Multilevel sequential2 monte carlo for bayesian inverse problems. *Journal of Computational Physics*, 368:154–178, 2018.

Trevelyan McKinley, Alex R Cook, and Robert Deardon. Inference in epidemic models without likelihoods. *The International Journal of Biostatistics*, 5(1), 2009.

Jamie Owen, Darren J Wilkinson, and Colin S Gillespie. Likelihood free inference for markov processes: a comparison. *Statistical Applications in Genetics and Molecular Biology*, 14(2):189–209, 2015.

Dennis Prangle. Adapting the abc distance function. *Bayesian Analysis*, 12(1):289–309, 2017.

Cristian Predescu, Mihaela Predescu, and Cristian V Ciobanu. The incomplete beta function law for parallel tempering sampling of classical canonical systems. *The Journal of Chemical Physics*, 120(9):4119–4128, 2004.

A. Raue, M. Schilling, J. Bachmann, A. Matteson, M. Schelke, D. Kaschek, S. Hug, C. Kreutz, B. D. Harms, F. J. Theis, U. Klingmüller, and J. Timmer. Lessons learned from quantitative dynamical modeling in systems biology. *PLoS ONE*, 8(9):e74335, Sept. 2013.

SA Sisson, Y Fan, and MA Beaumont. Overview of abc. *Handbook of Approximate Bayesian Computation*, pages 3–54, 2018.

Yuji Sugita, Akio Kitao, and Yuko Okamoto. Multidimensional replica-exchange method for free-energy calculations. *The Journal of Chemical Physics*, 113(15):6042–6051, 2000.

Tina Toni, David Welch, Natalja Strelkowa, Andreas Ipsen, and Michael PH Stumpf. Approximate bayesian computation scheme for parameter inference and model selection in dynamical systems. *Journal of the Royal Society Interface*, 6(31):187–202, 2008.

WD Vousden, Will M Farr, and Ilya Mandel. Dynamic temperature selection for parallel tempering in markov chain monte carlo simulations. *Monthly Notices of the Royal Astronomical Society*, 455(2):1919–1937, 2015.

Daniel Weindl, Yannik Schälte, Jan Hasenauer, Paul Stapor, Fabian Fröhlich, Elba Raimúndez Alvarez, and Charles Tapley Hoyt. Icb-dcm/petab: Petab v0.0.0a13, April 2019. URL https://doi.org/10.5281/zenodo.2630875.

Richard David Wilkinson. Approximate bayesian computation (abc) gives exact results under the assumption of model error. *Statistical Applications in Genetics and Molecular Biology*, 12(2):129–141, 2013.
